# The timing of cellular events: a stochastic vs deterministic perspective

**DOI:** 10.1101/2023.07.20.549956

**Authors:** Lucy Ham, Megan A. Coomer, Kaan Öcal, Ramon Grima, Michael P.H. Stumpf

**Affiliations:** School of BioSciences, University of Melbourne, Parkville VIC 3010, Australia; School of Mathematics and Statistics, University of Melbourne, Parkville VIC 3010, Australia; School of Informatics, University of Edinburgh, Edinburgh, EH8 9AB, United Kingdom; School of Biological Sciences, University of Edinburgh, Edinburgh, EH9 3JH, United Kingdom

**Keywords:** first-passage time, stochastic, deterministic, moments, event timing, threshold

## Abstract

Changes in cell state are driven by key molecular events whose timing can often be measured experimentally. Of particular interest is the time taken for the levels of RNA or protein molecules to reach a critical threshold defining the triggering of a cellular event. While this mean trigger time can be estimated by numerical integration of deterministic models, these ignore intrinsic noise and hence their predictions may be inaccurate. Here we study the differences between deterministic and stochastic model predictions for the mean trigger times using simple models of gene expression, post-transcriptional feedback control, and enzyme-mediated catalysis. By comparison of the two predictions, we show that when promoter switching is present there exists a transition from a parameter regime where deterministic models predict a longer trigger time than stochastic models to a regime where the opposite occurs. Furthermore, the ratio of the trigger times of the two models can be large, particularly for auto-regulatory genetic feedback loops. Our theory provides intuitive insight into the origin of these effects and shows that deterministic predictions for cellular event timing can be highly inaccurate when molecule numbers are within the range known for many cells.

## 1. Introduction

Cellular dynamics are not deterministic [1]. Observations of RNA or protein numbers in single cells show that their variation with time is stochastic [2, 3, 4]. This noise originates from biomolecular processes that are unaccounted for [5, 6], as well as from the inherent random timing of individual molecular events which manifests most clearly in low copy number conditions (intrinsic noise) [7, 8]. In spite of this randomness, cells orchestrate their processes with remarkable precision and robustness, likely because of mechanisms which either exploit or suppress intrinsic noise [9, 10].

Mathematical modelling based on the chemical master equation (CME), a probabilistic description of chemical reaction kinetics, has led to significant progress in understanding how the stationary and non-stationary distributions of the number of molecules in a single cell vary as a function of the rate parameter values [11, 12, 13, 14, 15]. Nevertheless these studies do not directly address a central question: when will the level of some molecule, first cross a critical threshold? It is known that regulatory proteins often need to reach critical threshold levels to trigger downstream processes [16, 17, 18, 19, 20, 21, 22, 23]. For example, in cellular development and differentiation, progression through different phases of the cell cycle only occurs once certain cyclin proteins surpass a prescribed threshold [24, 25]. Another example is p53-mediated cell apoptosis which is triggered when p53 proteins reach a critical value; below this value, cells enter the G1 arrest phase [26, 27] (Figure 1(A)). In the context of RNA, it has been shown that microRNAs (miRNAs) can create mRNA threshold levels, below which protein production is suppressed [28, 29]. From an experimental perspective, threshold crossing can however be understood more generally. For example, many events in cells can be timed by the turning on or off of a molecular fluorescent marker; since the observed molecular levels are of a continuous nature, a threshold is necessary to define a discrete event from these markers. Examples of processes that have been timed in this way include entry into competence in bacteria, different phases of lysis by phage, cell cycle phases, and loss or gain of pluripotency in stem cells (for a review see [30]).

**Figure 1:**
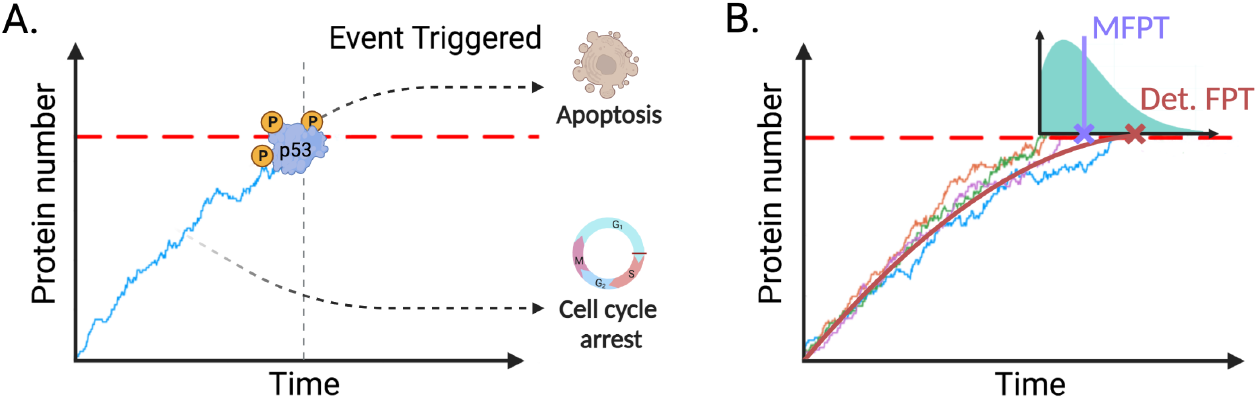
(A) Many processes rely upon regulatory proteins or molecules attaining critical threshold levels. Once this threshold is surpassed, a biological event is triggered. The time at which this occurs is known as the *first-passage time* (vertical grey dashed line). For example, upon DNA damage, the activation of p53 can elicit two potential cellular responses: cell-cycle arrest or apoptosis. Below a defined threshold of p53 expression, the cell undergoes cell-cycle arrest, whereas apoptosis is induced when the p53 expression surpasses a threshold. (B) The underlying process governing the regulatory molecule is stochastic and hence variability in the event timing is expected. This results in a distribution of first-passage times (shown in turquoise), which is conditional on the initial number of protein molecules in the system; here the initial protein number is 0. The mean first-passage time (MFPT; solid purple line and cross) can be compared with the deterministic FPT (red cross) which is the time to reach the target molecular number as estimated by integration of deterministic reaction rate equations (red solid line).

The time for a molecule number to reach a certain target value is a stochastic variable whose properties can be understood using first-passage time (FPT) theory [31]. Of particular importance is the mean first-passage time, the average time to reach a target value. Analytical expressions for the mean FPT have been derived for one variable models [32, 33], and systems with many states connected in a simple way [34, 35, 36]. For more complex models, as common in biology, the exact analytical approach is often unfeasible and thus numerical computation, using the Finite State Projection (FSP) method [37, 38], or Monte Carlo simulation using the Stochastic Simulation Algorithm (SSA) [39], is typically far more practical. Approximations, such as moment closure approaches, might be feasible too [40, 41]. Using one or more of these approaches, several studies have investigated the mean FPT (and higher-order moments and distributions of the FPT) as a function of the rate parameter values [42, 43, 44, 45, 46, 47, 48, 49, 13]. Despite these advances, biochemical systems have been, and are still, studied by means of deterministic ordinary differential equations due to their simplicity and ease of use [50]. Within this framework, it is also possible to estimate the time to reach a certain molecular number threshold by numerical integration of the differential equations. The difference between estimates using deterministic and stochastic modelling frameworks have not previously been systematically studied except in special cases (which we discuss below). This is an important question given the ubiquity of deterministic models of cellular biology and the fact that comparison of the two estimates would provide a direct means to study the role played by intrinsic noise in the timing of cellular events.

There are a few limits where the relationship of the deterministic and stochastic predictions for the mean time to reach a target molecular threshold are well understood. If for a given initial condition, the target molecule number is outside of the range predicted by the deterministic model in finite time then clearly the latter’s prediction for the mean time is undefined whereas a stochastic model will typically predict a finite value. An extreme example is given by extinction processes, where initially the number of molecules is larger than zero and the target value is zero, a value that the deterministic model only approaches asymptotically [51, 52]. If the target molecule number is within the range predicted by the deterministic model in finite time then, if intrinsic noise is very small (molecule numbers are quite large), the trajectories obtained from the SSA are close to the temporal variation of the mean molecule numbers predicted by the deterministic model [32, 53] and hence the stochastic and deterministic approaches will necessarily predict the same mean time to reach a threshold molecule number.

The relationship between deterministic and stochastic predictions of the mean time to reach a threshold value is, however, unclear when the molecule numbers are small—the typical case inside cells. Even if the mean molecule numbers of the deterministic and stochastic models are the same for all times (as for example is the case for systems with linear propensities [33]), when intrinsic noise is large, the trajectories of the SSA will not closely follow the deterministic prediction. For systems with nonlinear propensities (such as those with bimolecular reactions) the differences are likely even more pronounced because the mean molecule number predictions of the two models disagree which will also be reflected in the trajectories of the SSA [54]. Hence generally it is difficult to say if the mean time it takes for trajectories of the SSA to reach a designated target level (as measured by the mean FPT) is the same, larger or smaller than the time predicted by the deterministic model. An illustration comparing the two model predictions is shown in Figure 1(B).

We shed light on this problem by computing the difference in trigger time predictions of deterministic and stochastic models of a variety of biochemical processes with the constraint that the target molecular value is larger than zero and smaller than the steady-state mean molecular number predicted by deterministic models. The analysis enhances our understanding of the role played by intrinsic noise in cellular event timing.

## 2. Mathematical framework

We can formulate the question of interest as a FPT problem: for a continuous-time Markov process (CTMP) **x**(*t*), we are interested in the time that it takes **x**(*t*) to first arrive at some subset **Y** of the state-space, given the system is started from **n**∉**Y**. In other words, we are interested in the distribution of first-passage times,

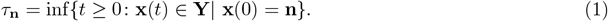

### 2.1. Moments of the FPT distribution

We derive moments of the FPT distribution for any CTMP **x**(*t*), providing a fresh intuitive proof of a result that has been presented in the literature [32, 45]. Fix an initial state **n** ∉ **Y** and let *τ*_**nz**_ be the waiting time to reach the absorbing state **Y** from **n** given that **z** is the next state visited. The random variable *τ*_**nz**_ can be decomposed as the sum of two independent random variables,

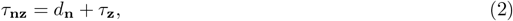

where *d*_**n**_ is the time until the first jump (the exponentially distributed dwelling time) and *τ*_**z**_ is the waiting time to reach the absorbing set **Y** from the state **z**. Let **A** denote the state transition matrix for **x**(*t*) (Supplementary Material, Section 1.1). The expected time until the next jump is −1/*A*_**nn**_, and the probability of transitioning to state **z** equals −*A*_**zn**_/*A*_**nn**_, where here *A*_*ij*_ is the *ij*^th^ element of **A**. For any k≥1, we can compute the k^th^ raw moment of the waiting time *τ*_**n**_ to reach **Y** from **n** as follows. By the Law of Total Expectation, we have that

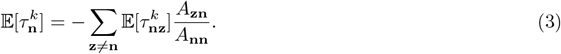

Using (2), the Binomial Theorem, and the independence of *d*_**n**_ from *τ*_**z**_, it follows that,

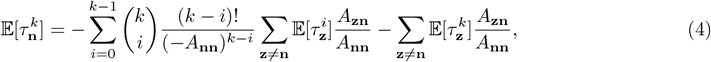

for **n** ∉ **Y**, where here we have also used the fact that *d*_*n*_ is exponentially distributed with rate −*A*_**nn**_. Now moving the rightmost sum to the left and multiplying both sides of (4) through by *A*_**nn**_, it follows that

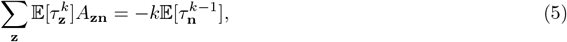

where here we have used (3) from right to left with *k* − 1 in place of *k*. Recalling that *τ*_**z**_ ≡ 0 for **z** ∈ **Y** and that **n** ∉ **Y**, we can write (5) for all initial states **z** ∉ **Y** concisely as

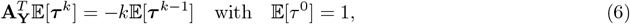

where 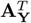 denotes the matrix **A**^*T*^ with rows and columns corresponding to **Y** removed. Solving this (matrix) recurrence relation gives,

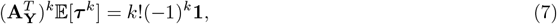

which agrees with the result that can be obtained from the Backward Chemical Master Equation (Supplementary Material, Section 2.1). We use a modification of the FSP algorithm to compute mean FPTs (MFPTs) via (7) (Supplementary Material, Section 1.2). Our approach allows us to simultaneously compute mean FPTs, for *all* initial molecule numbers **n** ∉ **Y**, significantly reducing computation time in comparison to Monte Carlo simulations; refer to Section 5 of the Supplementary Material, where we compare CPU times for three models of biological relevance, including enzyme kinetic examples and a compartmental model of disease spread. We remark that the Forward CME can be used to compute FPT distributions by way of a modified FSP, however requires recomputation for each initial state [38]. Note that (7) can also be solved analytically in some cases (Sections 3 and 4 of the Supplementary Material).

## 3. Stochastic vs deterministic calculations

In the following sections, we will refer to the time that the deterministic process first reaches a given threshold as the *deterministic FPT*.

### 3.1. The bursty birth-death process

We consider the bursty birth-death process with fixed burst size *r*,

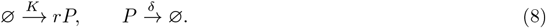

See Figure 2(A) for an illustration of this model. It is assumed there is the continuous production of a molecular species *P* in bursts of fixed size *r*, according to a Poisson process at constant rate *K*. The degradation process captures both active degradation as well as dilution due to cell division, and is modelled as a first-order Poisson process with rate *δ*. Throughout the remainder of the paper, we rescale time as *τ* = *δ* · *t*, and assume that all model parameters have been scaled by the degradation rate *δ*, so that *δ* = 1. Hence both time and the rate parameters are non-dimensional. The bursty birth-death process serves as a crude model for bursty expression of either mRNA or protein due to transcription or translation; we do not here account for the fact that the burst size in these cases is not fixed, but is distributed according to a geometric distribution [11]. For specificity, in what follows we shall refer to *P* as protein.

**Figure 2:**
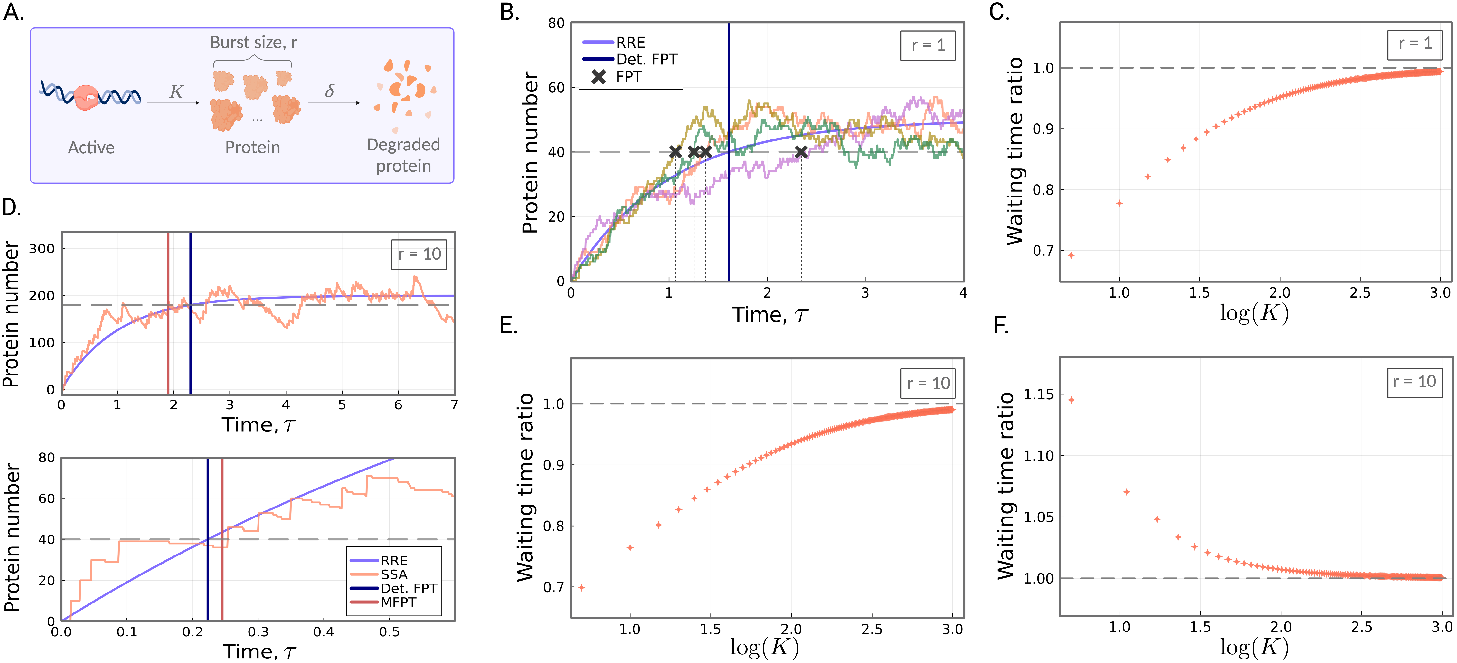
Stochastic *vs*. deterministic waiting times to reach a target protein number for the bursty birth-death process. (A) An illustration of the bursty birth-death process (as given by the reaction scheme in (8) with burst size *r*≥1). (B) Example trajectories of the simple birth-death process (multi-coloured) as simulated by the SSA. The FPTs of the trajectories are shown as the dark grey crosses, while the deterministic FPT is shown as the blue vertical line. Here *r* = 1, *ρ* = 0.8, *K* = 50 and *δ* = 1, so that the target *N* = 40 (grey dashed line). (C) The ratio of the stochastic MFPT to the deterministic FPT as a function of increasing *K*. The stochastic mean waiting time is computed according to (9), and the deterministic waiting time is computed according to (11). Model parameters are *ρ* = 0.8, *K* varies from 2 to 10^3^, and *δ* = 1. (D) (top) Waiting times when target *N* is equal to a large proportion (here *ρ* = 0.9) of the steady-state mean. A representative time series of the bursty birth-death process is shown in orange, and the solution of the reaction rate equation in purple. Here *r* = 10, *K* = 20 and *δ* = 1, so that the target *N* = 180 (shown as the grey dashed line). The stochastic MFPT (red vertical line) is smaller the deterministic FPT (blue). (bottom) Parameters are the same as above except now *ρ* = 0.2 and the target is *N* = 40. (E) The ratio of the stochastic MFPT to the deterministic FPT as a function of increasing *K*, for a high threshold value. The number of initial proteins is 0, *ρ* = 0.9, *r* = 10, *K* varies from 2 to 10^3^, and *δ* = 1. (F) Same as (E), except now *ρ* = 0.2. Stochastic waiting times are computed according to our FSP approach (refer to Section 1.2 of the Supplementary Material), and deterministic waiting times are computed analytically.

#### Case 1: the simple birth-death process

When *r* = 1, the model given in reaction scheme (8) coincides with a simple birth-death process. We are interested in the mean waiting time for the birth-death process to reach a fixed protein number *N*, given that the system is started from *n* initial proteins. Simulations of the simple birth-death process suggest the MFPT is smaller than the deterministic FPT (see Figure 2(B), but note that only a small sample of trajectories are shown here for visualisation purposes). We now prove this analytically. To begin, we find an analytical solution for the MFPT of the stochastic description of the system, as given by the CME (see Eq. (3) of the Supplementary Material).

Starting from (7), it can be shown (Supplementary Material, Section 3), that the expected waiting time to reach *N* proteins given the system is started from *n* ≤ *N*, is given by,

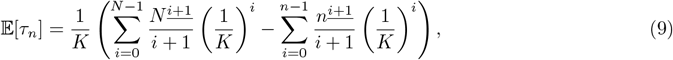

where for real number *x* and positive integer *m*, the notation *x*^*<u>m</u>*^ abbreviates *x*(*x* −1) … (*x* −(*m*−1)), the falling factorial of *x*. In what follows for simplicity we shall set n = 0. The deterministic description of the system is given by the reaction rate equation *d*_*τ*_ *X* = *K* − *X*, with *X*(0) = 0, and solving for *X*(*τ*) yields *X*(*τ*) = K(1 − *e*^*−τ*^). Solving for *τ* then gives,

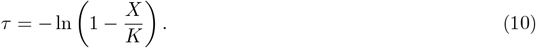

Consider some proportion 0 < *ρ* < 1, and consider the expected waiting time to reach *ρK*, that is, some proportion *ρ* of the steady-state mean. Then (10) simplifies to

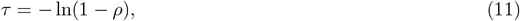

which we note is independent of *K*, and where the requirement of *ρ* < 1 enables this to be well defined. Now let us consider (9) for *N* = ⌊*ρK*⌋. For notational simplicity only, we assume that *ρK* is an integer, and again observe that the numerator *N* ^<u>*i*+1</u>^ is bounded above by (*ρK*)^*i*+1^. Replacing this in (9) gives

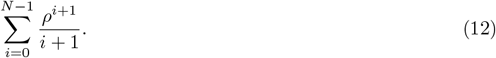

Recognising the Taylor expansion of −ln(1 −*ρ*) around 0 (for *ρ* < 1), we find that (12) is the truncated Taylor expansion of (11). Thus, as each term of the series is strictly positive, it follows that for any proportion *ρ* of the steady-state mean, the deterministic waiting time is a strict upper bound for the mean waiting times. An example is given in Figure 2(C) for *ρ* = 0.8. Here we show how the ratio of the stochastic mean waiting time to the deterministic waiting time of the simple birth-death process scales with increasing K. Note that since *δ* = *r* = 1, *K* is equal to the steady-state mean number of molecules.

#### *Case 2: burst size* r > 1

We now consider the case where protein is translated in bursts of size *r* > 1. Again, we are interested in the waiting time for the bursty system to reach some fixed target N given that the system starts with *n* = 0 initial proteins. In Figure 2(D) (top), we consider the case of a high threshold value; here the target *N* is equal to a large proportion (90%) of the steady-state mean, *Kr*. Noting that the deterministic model will not reach the steady-state mean in finite time (unlike the stochastic model), we expect a faster MFPT as predicted by the stochastic model than that predicted by the deterministic model. Indeed, we see that the stochastic MFPT to reach *N* (red vertical line) is again smaller than the deterministic FPT (blue vertical line). In Figure 2(E) (left), we show how the ratio of the MFPT to the deterministic FPT scales with increasing *K*, for a high threshold. The differences are most pronounced for systems with a smaller steady-state mean number molecules, with waiting times eventually converging in the limit of large mean molecule number.

To build intuition for the opposite case, that is, targets that are a small proportion of the steady-state mean, we again look to the extremes. If the target *N* is chosen such that *N* < *r*, that is, strictly below the burst size, then the expected waiting time to surpass this in the stochastic case is simply the expected waiting time to the first event, which is 1/*K*; this is independent of *N*. But as *N* tends to 0, the deterministic solution also tends to 0, so that a small enough choice of *N* will place it faster than 1/*K*; the precise boundary for this can be derived as *N* < *Kr*(1 −*e*^*−*1*/K*^) ≈*r*. Note that this condition cannot hold for the simple-birth death process as *r* = 1 and *N* is discrete. In Figure 2(D) (bottom), we see that this difference extends to targets that are much higher than the burst size; here *N* = 40 and *r* = 10. The waiting time as predicted by the deterministic solution (blue vertical line) is indeed smaller than the MFPT to reach N (red vertical line). Comparing the trajectories in Figure 2(D) top and bottom, we see that the deterministic solution is faster than the stochastic solution when the target threshold is low, in agreement with our theoretical argument above. In Figure 2(F), we show how the ratio of the stochastic MFPT to the deterministic FPT scales with increasing *K*, for a low threshold value. The differences are most pronounced when the steady-state mean molecule numbers are small.

### 3.2. The telegraph model

Next we investigate whether or not the same transition in the ratio of the deterministic to stochastic waiting times from below to above 1 is possible using a more realistic model of gene expression. We consider the two-state telegraph model of gene expression [55], which has been widely employed in the literature to model bursty gene expression in eukaryotic cells [56, 57]; a schematic is given in Figure 3(A). The effective reactions for the telegraph model are given by,

**Figure 3:**
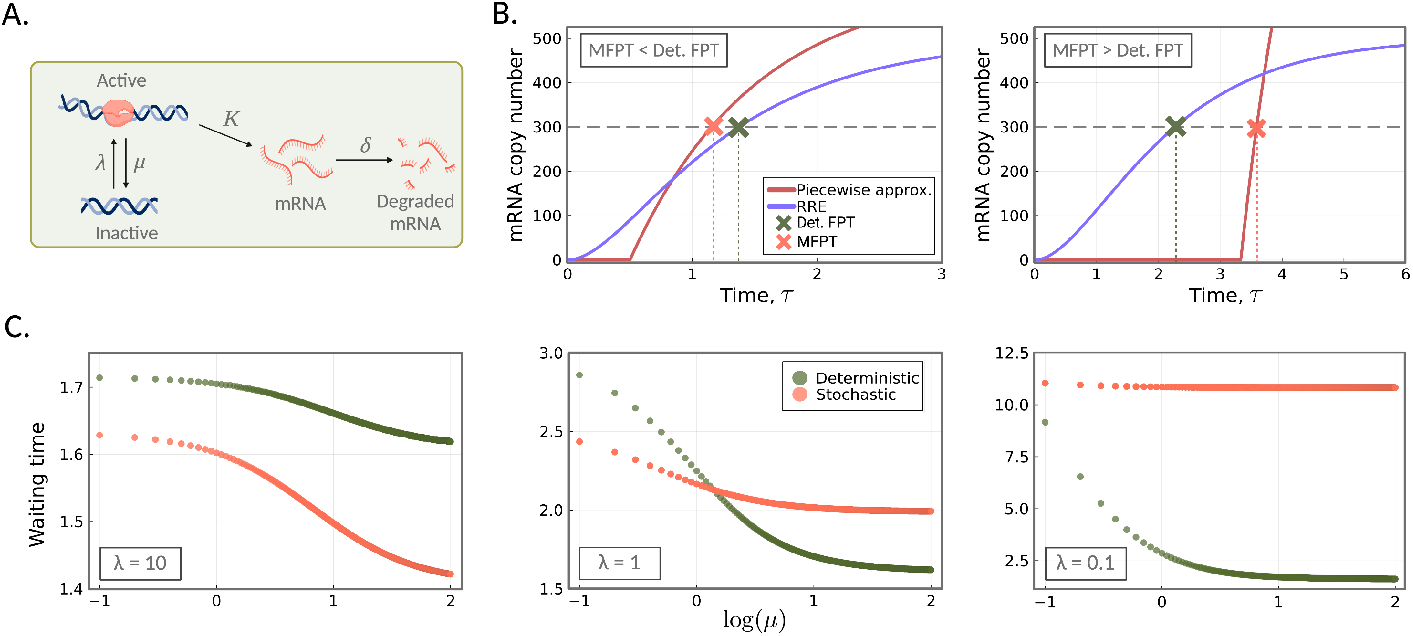
Stochastic *vs*. deterministic waiting times to reach a target mRNA value for the telegraph model. (A) A schematic of the telegraph model as given by the reaction scheme in (13). (B) Piecewise-deterministic approximation of the stochastic model (14) versus the deterministic model. (left) For large *λ* values, the approximate MFPT (orange cross) of the telegraph process to reach target *N* (grey dashed line) is smaller than the deterministic FPT (green cross). The deterministic reaction rate equation is shown in purple. Model parameters are *λ* = 2, *μ* = 0.5, *K* = 625, so that the steady-state mean mRNA copy number is 500. We also have that *ρ* = 0.6, so that the target *N* = 300. (right) For small values of *λ*, the approximate MPFT of the stochastic model (orange cross) is larger than that of the deterministic model (green cross). Model parameters used are *λ* = 0.3, *μ* = 0.5, *K* = 1333, and *ρ* = 0.6. (C) FPTs under deterministic (green) and stochastic modelling (orange; computed using FSP) as a function of the switching off rate *μ*, for three different *λ* values: *λ* = 10 (left), *λ* = 1 (middle), *λ* = 0.1 (right). In each plot, we vary *μ* from 0 to 100, while keeping the steady-state mean mRNA number fixed at 100; this involves varying *K* accordingly. The target *N* is set to 80% of the steady-state mean.

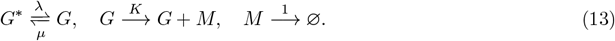

In this model, the gene switches probabilistically between an inactive state *G*^*∗*^ and active state *G* from which mRNA molecules (*M*) are synthesised; degradation of mRNA occurs independently of the promoter state.

Again, we are interested in examining waiting times of the telegraph system to reach some target 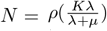, that is, some proportion 0 < *ρ* < 1 of the steady-state mean number of molecules. In contrast to the model considered in the previous section, the gene can now experience periods of inactivity in which no transcription occurs. If the telegraph system is initialised in the inactive state *G*^*∗*^, then the waiting time to reach *N* starting from 0 molecules is *at least* the expected waiting time for the system to transition to the active state (the first event), which is equal to 1/λ. Thus, 1/λ is a strict lower bound for the mean first-passage time of the system to reach the target N.

If we assume that mRNA is abundant enough so that conditional on the gene state the dynamics are deterministic, then the mRNA dynamics, from *τ* = 0 to *τ* = *τ*_off_, are approximated by,

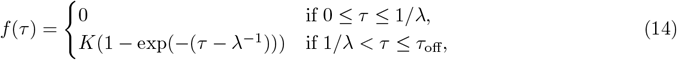

where *τ*_off_ = (1/λ) + (1/*μ*) (mean total time for the gene to turn on and then off). Note that the first line describes no expression while the gene is off whilst the second describes the switching on of gene expression. In particular, for times close to the switching on time, we have that *f*(*τ*) ≈*K*(*τ*− λ^*−*1^), implying there is a sudden linear increase in expression with a slope that is dependent on the mRNA synthesis rate *K*. In contrast, the deterministic model solution is non-zero for all times *t* > 0 because in this case both the gene and mRNA are both treated in a continuous manner. Thus, due to the immediate initiation of mRNA production in the deterministic model, it consistently maintains a competitive edge over the stochastic model in terms of reaching the target first. However, the stochastic model possesses a notable advantage: upon gene activation, the initiation of mRNA production progresses considerably faster compared to the deterministic model. Because of these two opposing properties, we expect the MFPT to be less than the deterministic FPT if the activation rate is sufficiently large (Figure 3(B) left) and the opposite if the activation rate is quite small (Figure 3(B) right).

These two scenarios are corroborated by computing the stochastic MFPTs via our FSP approach (Figure 3(C) left and right, respectively). Note that in these plots we are varying the switching off rate *μ* and synthesis rate *K* such that the steady-state mean mRNA is unchanged. While the variation of *K* changes the sharpness of the mRNA response when a gene switches on, its effect on the difference between the MFPT and its deterministic equivalent is secondary to the influence of the switching on rate *λ*, provided this is very large or very small. However, when λ is moderately large then the value of *K* becomes the determining factor (Figure 3(C) middle). We remark that while the piecewise function (14) well approximates the mRNA for large and small λ values, for intermediate values the theory is unable to account for the fact that a proportion of trajectories will not reach the target on the time interval 0 ≤*τ* ≤*τ*_off_ – hence this approximation cannot be used to accurately predict the value of *μ* at which there is a transition in the waiting time ratio from above to below 1 in Figure 3(C) middle.

Furthermore, note that transition in the waiting time ratio seen in the telegraph model is broadly speaking similar in character to that in the simpler bursty birth-death model. Large λ means the gene is mostly on and hence this is similar to constitutive expression (the the birth-death model with *r* = 1) for which we proved the deterministic FPT to be an upperbound for the MFPT of the stochastic model (Figure 2(C)). Small λ means the gene is mostly off and if the synthesis rate is large enough this implies bursty (*r* > 1) expression for which the MFPT can be larger than its deterministic counterpart (Figure 2(F)). Given the qualitative similarities in the predictions of the two models, it is likely the form of the burst size distribution (a delta function for the bursty birth-death model and a geometric distribution for the telegraph model in certain limits [12]) is not important to the existence of the observed transition in the waiting time ratio.

### 3.3. An autoregulatory feedback loop

Next we examine an autoregulatory feedback loop in which a protein produced by a gene either enhances or suppresses its own expression. This motif is commonly encountered in biology. For example, in *E. coli* negative autoregulation appears in over 40% of known transcription factors. A number of studies have used stochastic models to investigate optimal feedback mechanisms to minimise the variability in FPTs [44, 58, 59, 46, 60, 61]. The reaction scheme for the feedback loop that we employ is given, by [62],

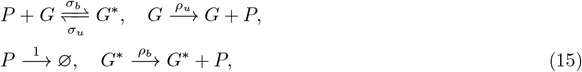

where *G*^*∗*^ and *G* represent the bound and unbound states respectively, and *P* represents protein. The parameters *σ*_*u*_ and *σ*_*b*_ are the unbinding and binding rates respectively, and *δ* is the protein degradation rate. The rate of protein production depends on the gene state and is given by *ρ*_*b*_ and *ρ*_*u*_ in the bound and unbound states, respectively. When *ρ*_*b*_ > *ρ*_*u*_ the feedback is positive since an increase in the protein number will cause the gene to switch more often to the *G*^*∗*^ state in which the production rate is high. Similarly when *ρ*_*b*_ < *ρ*_*u*_ the feedback is negative. We illustrate the positive and negative feedback loops in Figure 4(A) left and right panels, respectively. More realistic models have been devised, for example [63] describes a bursty version of (15) to account for translational bursting; however these details are unlikely to change the qualitative properties of the waiting time results that we will describe next.

**Figure 4:**
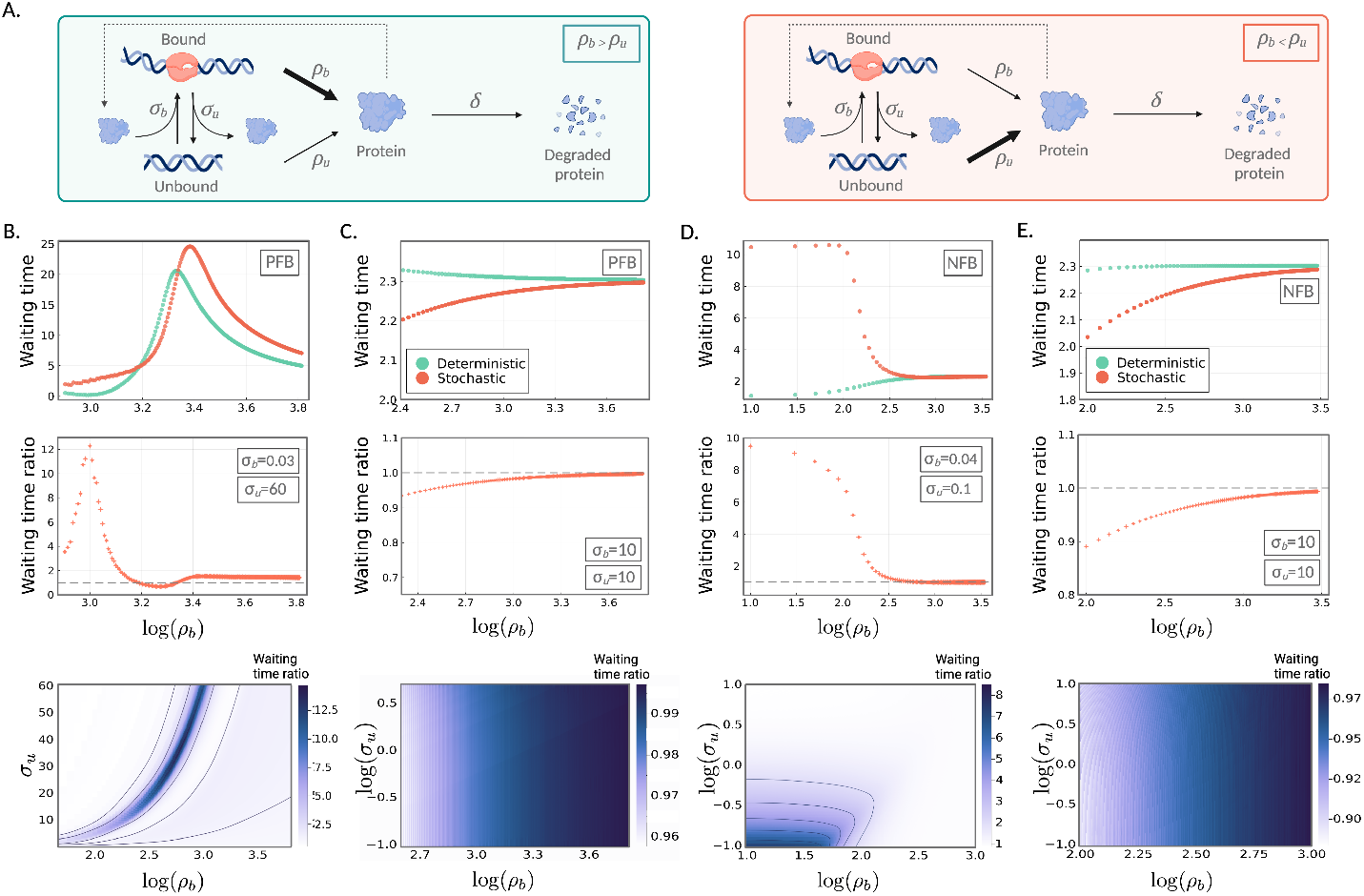
Stochastic *vs*. deterministic waiting times to achieve a target protein value for auto-regulatory feedback models (A) (left) An illustration of the feedback loop (15) for positive feedback (PFB) (i.e., *ρ*_*u*_ *< ρ*_*b*_), and (right) negative feedback (NFB) (i.e., *ρ*_*u*_ *> ρ*_*b*_). (B-E) Top panel: MFPT of the stochastic system (orange) to reach 90% of the steady-state mean, given the system is started from 0 protein molecules, for increasing *ρ*_*b*_. The corresponding deterministic waiting times are shown in green. Middle panel: the corresponding ratio of the stochastic waiting time to the deterministic waiting time, as a function of *ρ*_*b*_. Bottom panel: ratio of the MFPT of the stochastic system to the deterministic FPT, to reach 90% of the steady-state mean, given the system is started from 0 protein molecules, as a function of *ρ*_*b*_ and *σ*_*u*_. All parameters along with the mean molecule numbers are given in Table 1 in the Supplementary Material.

In Figure 4(B)&(C) (vertical panels), we consider the case of positive feedback (i.e., when *ρ*_*u*_ < *ρ*_*b*_). For two different *σ*_*b*_ values, we show the MFPT of the stochastic system (orange) to reach 90% of the steadystate mean, given the system is started from 0 protein molecules, for increasing *ρ*_*b*_; this can be thought of as increasing the strength of the positive feedback. The corresponding deterministic waiting times are displayed in green. For low values of the binding rate *σ*_*b*_ (Figure 4(B)), we see a rapid rise and fall in the FPTs (for both the stochastic and deterministic models), as we increase the strength of positive feedback. This initial rise in the waiting time may seem somewhat counterintuitive, but can be understood in terms of the corresponding increase in the steady-state mean. As the transcription rate *ρ*_*b*_ rises, so too does does the steady-state mean, and hence the target *N*. Because *σ*_*b*_ is very low, binding is a rare event and short lived, as *σ*_*u*_ is large. This means that in a typical trajectory, most of the time towards reaching the target is spent in the unbound state with constant transcription rate *ρ*_*u*_, attempting to reach a rising target. For high values of *ρ*_*b*_, the level of transcription is sufficiently high that even the short window of relatively rare bound behaviour provides significant progress towards achieving the target molecule number. For high *σ*_*b*_ values (Figure 4(C)), the effective binding rate, that is *σ*_*b*_⟨*P*⟩, quickly becomes enormously higher than the unbinding rate *σ*_*u*_, so that the system is very close to constitutive, and the deterministic FPTs bound from above the stochastic MFPTs. The corresponding waiting-time ratio plots also reveal that this ratio can be quite large when it is above 1 (an order of magnitude) – this is a considerably larger effect than seen for the bursty birth-death and telegraph models, likely because feedback models have a bimolecular reaction which enhances the differences between the SSA trajectories and the deterministic models [54]. Note that the largest differences in the deterministic and stochastic waiting times occur within the range of tens to hundreds of molecules (here the mean number of molecules is approximately between 11.68 and 370.91). This is within the range known for many cells. For example, in E. Coli, copy numbers of transcription factors are quite low, occurring between 1–1000 per cell [64, 65]. The heatmaps in Figure 4(B)&(C) (bottom plots) show that the behaviour in the top and middle plots are qualitatively representative across a broad range of *σ*_*u*_ values.

In the case of negative feedback (i.e., when *ρ*_*b*_ < *ρ*_*u*_), increasing *ρ*_*b*_ now leads to a decrease in the strength of negative feedback, and hence again a rise in the overall mean steady-state protein number. Thus, the overall trends are broadly similar to the positive feedback case. For a low *σ*_*b*_ value (see Figure 4(D) with *σ*_*b*_ = 0.04), we intially see a very slight increase in the MFPT in the region of low *ρ*_*b*_. This is followed by a rapid decline in the MFPT, with high values of *ρ*_*b*_ (that is, approaching equality with *ρ*_*u*_), becoming close to a constitutive model. We note that again the largest differences in the stochastic and deterministic waiting times occur within the range of tens to hundreds of molecules (between approximately 44.36 and 317.1). For high *σ*_*b*_ (Figure 4(E) with *σ*_*b*_ = 10), the effective binding rate is significantly higher than *σ*_*u*_, so that the system is close to being constitutive with protein production rate at *ρ*_*b*_. Hence the stochastic MFPTs are bounded above by the deterministic FPTs; refer to Figure 4(E) (top and middle). The heatmaps in Figure 4(D)&(E)(bottom) again show that the parameter sets in (D)&(E) (top and middle) are qualitatively representative across a broad range of *σ*_*u*_ values.

## 4. Summary and Discussion

In this article, we have investigated the difference between deterministic and stochastic model predictions for the timing of cellular events when the target molecular value can be reached by the deterministic model in finite time. While differences between the mean molecule number predictions of these models have previously been extensively investigated [54, 66], differences in their predictions of the mean time to trigger a cellular event have not.

Our analysis elucidated a number of interesting results which we now summarise. For the simple birthdeath process, where molecules are produced and destroyed one at a time (a model for constitutive gene expression if the molecules are assumed to be mRNA [67]), we have proven that the deterministic model prediction provides a strict upper bound for the mean trigger time predicted by the stochastic model. In other words, intrinsic noise leads to a reduction of the trigger time – this effect becomes more appreciable as the mean steady-state molecule number decreases. For a bursty birth-death process, where molecules are produced in batches of size *r* > 1 and destroyed one at a time, as the target molecular value is decreased, there is a switch from a regime where intrinsic noise reduces the mean target time to one where it lengthens it (Figure 2(D)). Alternatively, the same transition can be observed if the target value is fixed, but the burst size r is increased. Since the expression of many genes is bursty (of transcriptional or translational origin), this simple model suggests that the transition may exist in more realistic models of gene expression. Indeed, we showed that the same transition occurs in the two-state telegraph model of gene expression [55, 67], which is capable of explaining both constitutive, bursty, and intermediate behaviour, and in a model of an autoregulatory genetic feedback loop, a common motif in nature. For the telegraph model, we showed that this transition can be explained in an intuitive mechanistic way by a simple quantitative description of the early-time kinetics using a piecewise-deterministic model of gene expression. For feedback loops, we additionally observed large ratios of the deterministic and stochastic mean target times compared to previous models, likely because these loops have a bimolecular reaction between protein and the gene. The effect of such reactions on the differences between the deterministic and stochastic mean molecule numbers are well known to increase with decreasing system size (volume of the system), and with increasing proximity to the point in parameter space where a system switches from stable to an unstable dynamical behaviour [54]. In fact, our analysis of the waiting times for the substrate to reach a certain target level in an enzyme-substrate reaction (Fig. S6) showed that the ratio of deterministic and stochastic mean waiting times also increased with these two system properties.

We note that our main analytical and numerical results (the transitions in the waiting time ratio) are not directly comparable with those of other FPT studies [42, 43, 44, 45, 46, 47, 48, 13]. This is because the focus of these studies was to understand FPT statistics as a function of the rate parameters, using either a stochastic model or deterministic approximation. From a computational perspective, we note that while other studies have also used FSP to calculate first-passage time statistics [38], our modification of the FSP for computing the statistics via (7) is advantageous because it does not require recomputation for each initial state. As we show in the SI, the method also leads to large computational savings compared to estimation of the FPT statistics using the SSA. Concluding, our study shows that intrinsic noise has non-trivial effects on the timing of cellular events and that simple models can provide intuitive insights into the microscopic origins of these effects.

## Acknowledgments

We gratefully acknowledge support from the members of the *Theoretical Systems Biology Group* at the University of Melbourne. L.H. and M.P.H.S. are supported by the University of Melbourne DVCR fund, M.A.C by the University of Melbourne graduate research scholarship, K.Ö. by the EPSRC Centre for Doctoral Training in Data Science (EPSRC grant EP/L016427/1) and R.G. by the Leverhulme Trust grant (Grant No. RPG-2020-327).

