## Supplementary Material for "The timing of cellular events: a stochastic vs deterministic perspective"

### Supplementary material for
### The timing of cellular events: a stochastic vs deterministic perspective

Lucy Ham[a,b,1], Megan A. Coomer[b,1], Kaan Öcal[c,d], Ramon Grima[d], Michael P.H. Stumpf[a,b,*]

[a]School of BioSciences, University of Melbourne, Parkville VIC 3010, Australia
[b]School of Mathematics and Statistics, University of Melbourne, Parkville VIC 3010, Australia
[c]School of Informatics, University of Edinburgh, Edinburgh, EH8 9AB, United Kingdom
[d]School of Biological Sciences, University of Edinburgh, Edinburgh, EH9 3JH, United Kingdom

---

**Abstract**

Here we provide details and proofs of the results given in *The timing of cellular events: a stochastic vs deterministic perspective.*

---

#### 1. Methods

##### *1.1. Stochastic Reaction Networks*

Consider a well-stirred mixture consisting of $N$ chemical species $S_1, \ldots, S_N$ that interact through $M$ chemical reactions $R_1, \ldots, R_M$,

$$R_j \equiv \sum_{i=1}^{N} s_{ij} x_i \xrightarrow{c_j} \sum_{i=1}^{N} r_{ij} x_i, \quad j \in \{1, \ldots, M\}, \tag{1}$$

where $x_i$ denotes the number of $S_i$ molecules in the system at time $t$, $s_{ij}$ and $r_{ij}$ are integers, and $c_j$ is the rate constant of reaction $R_j$ with units of inverse time. Throughout, we will let $\mathbf{x}(t) = (x_1(t), \ldots, x_N(t))$ represent the state of the system at time $t$. Each reaction $R_j$ has an associated *propensity function* $a_j$ given by,

$$a_j(\mathbf{x}) = c_j h_j(\mathbf{x}), \tag{2}$$

where $h_j(\mathbf{x})$ is defined to be the number of distinct combinations of $R_j$-reactant molecules available in the state $\mathbf{x}$. The *state-change*, or *stoichiometry vector* $\mathbf{v}_j$ is defined to be the vector $(v_{1j}, \ldots, v_{Nj})$ whose $i$th component is given by $v_{ij} = r_{ij} - s_{ij}$, for $i \in \{1, \ldots, N\}$ and $j \in \{1, \ldots, M\}$. The process $\mathbf{x}(t)$ is a continuous-time Markov process, and the time evolution of the joint probability distribution of the molecule numbers is described by the Chemical Master Equation (CME),

$$d_t\mathbf{P} = \mathbf{A}\mathbf{P}, \tag{3}$$

where $\mathbf{P} := [P(\mathbf{x}_1), P(\mathbf{x}_2), \ldots]^T$ and $\mathbf{A}$ is the state transition matrix with the following structure [1],

$$A_{ik} := \begin{cases} -\sum_{j=1}^{M} a_j(\mathbf{x}), & \text{for } i = k \\ a_j(\mathbf{x}_i), & \text{for all } k \text{ such that } x_k = x_i + \mathbf{v}_j \\ 0, & \text{Otherwise.} \end{cases} \tag{4}$$

[1]These authors contributed equally to this work

*1.2. Finite State Projection for the modified CME*

Numerical computation of first-passage time (FPT) distributions can be performed using an adaptation of the Finite State Projection (FSP) [2]. Here the state space of the stochastic reaction network, which is usually infinite, is reduced to a finite subset consisting of the most relevant states. This converts the CME into a finite linear system of equations that can be solved efficiently on a computer.

In the standard formulation of the FSP, we select a finite set $\mathbf{Z}$ of states, which we assume to include the initial set, as well as the target set $\mathbf{Y}$. The terms in the CME corresponding to any state outside of $\mathbf{Z}$ are further assumed to vanish. Mathematically, this corresponds to solving the exact CME for a modified reaction network, wherein we combine all states outside of $\mathbf{Z}$ into a single state $\mathbf{Z}^c$ and remove all transitions from $\mathbf{Z}^c$ to $\mathbf{Z}$. Comparing this with the matrix $\mathbf{A}_{\mathbf{Y}}^T$, which contains all states excluding those in $\mathbf{Y}$, we see that applying the FSP to the construction of the matrix $\mathbf{A}_{\mathbf{Y}}$ eliminates all states that are either outside of the truncation, or in the absorbing set $\mathbf{Y}$. That is, we compute exactly the FPT distribution until the system either enters $\mathbf{Y}$ or leaves the truncated state space.

As a result, numerically estimating FPTs using the FSP will always underestimate the true first passage times, since the system leaving the truncated space is treated the same as the system entering the absorbing set $\mathbf{Y}$. To minimise the approximation error, therefore, one should choose the truncation $\mathbf{Z}$ such that the system is unlikely to leave $\mathbf{Z}$ before entering the target set $\mathbf{Y}$.

To compute FPTs numerically, we use the `FiniteStateProjection.jl` package [3] to construct the matrix $\mathbf{A}_{\mathbf{Y}}$, and solve the system of equations (15) (alternatively, Eq. (7) of the main text) using the standard sparse solvers provided in Julia.

#### 2. Computing moments of the FPT distribution

In the main text, we derive moments of the FPT distribution for any continuous-time Markov process $\mathbf{x}(t)$. We here illustrate our approach by way of a simple example. Consider a simple birth-death process with transcription rate $K$ and degradation rate $\delta$,

$$\varnothing \underset{\delta}{\overset{K}{\rightleftharpoons}} P. \tag{5}$$

Here $P$ represents protein. An illustration of the simple birth-death process is given in Figure 1(A). The state of the system, $S$, is the set of non-negative integers, $S = \{0, 1, 2, \dots\}$. Fix some target protein number $N \in S$, and suppose we are interested in the average time that it takes for the birth-death process to reach $N$ given that the system is started in state $n \leq N$. In other words, we are interested in finding,

$$\mathbb{E}[\tau_n] = \mathbb{E}[\inf\{t \geq 0 \text{ such that } X(t) = N\}|\ X(0) = n].$$

To find $\mathbb{E}[\tau_n]$, we begin by finding an expression for the average time that it takes to reach $N$ from $n$, conditional on the next state, which we will denote by $n'$. Letting $D_n := n\delta$ and $R_n := n\delta + K$, we may observe some simple linear relationships between neighbouring values of $\mathbb{E}[\tau_n]$. From a value of $n$, there are just two neighbouring states $n' = n+1$ (birth) or $n' = n-1$ (death). And these occur with birth and death probabilities $K/R_n$ and $D_n/R_n$, respectively. The time that it takes to get from $n$ to $N$ is the time that it takes to get from $n$ to $n'$ plus the time that it takes to get from $n'$ to $N$. Hence the expectation of the time taken to get to $N$ from state $n$, given the system transitions to $n'$, is $R_n^{-1} + \mathbb{E}[\tau_{n'}]$. It then follows from the Law of Total Expectation that,

$$\mathbb{E}[\tau_n] = R_n^{-1}\left[D_n(R_n^{-1} + \mathbb{E}[\tau_{n-1}]) + K(R_n^{-1} + \mathbb{E}[\tau_{n+1}])\right].$$

Rearranging, we obtain the tridiagonal recurrence relation,

$$R_n\mathbb{E}[\tau_n] - D_n\mathbb{E}[\tau_{n-1}] - K\mathbb{E}[\tau_{n+1}] = 1, \tag{6}$$

with $\mathbb{E}[\tau_N] = 0$. We impose $\tau_n = 0$ for $n < 0$ so that the recurrence relation is valid for $n \in \{0, 1, \dots, N-1\}$. Taking this set as our state space $S_N$ can write (6) in matrix form as,

$$\mathbf{A}\mathbb{E}(\boldsymbol{\tau}) = \mathbf{1}, \tag{7}$$

where $\mathbf{A}$ is an $N \times N$ tridiagonal matrix defined by,

$$\mathbf{A}_{ij} := \begin{cases} R_i, & \text{for } i = j \\ -D_i, & \text{for } j = i+1 \\ -K, & \text{for } j = i-1 \\ 0, & \text{Otherwise.} \end{cases} \tag{8}$$

Here $\mathbf{1} = [1, 1, \dots, 1]^T$ is a vector of $N$ 1's, and $\mathbb{E}(\boldsymbol{\tau}) = [\mathbb{E}[\tau_0], \dots, \mathbb{E}[\tau_{N-1}]]^T$ is the unknown vector of mean first passage times. (7) constitutes a finite system of linear equations which can solved numerically, which is the basis of our modified FSP approach (see subsection 1.2 above), or analytically in some cases (see Section 3 below).

##### *2.1. Relationship to the Backward Master Equation*

In the main text, we derive moments of the FPT distribution of arbitrary order (Eq. (7) of the main text). We here demonstrate how our result agrees with that obtained from the Backwards Chemical Master Equation (BCME). We begin by observing the standard relationships between the FPT distribution and the cumulative and survival distributions,

$$F_{\mathbf{n}}(t) = \int_0^t f_{\mathbf{n}}(s)\ ds. \tag{9}$$

We thus have that,

$$f_{\mathbf{n}}(t) = -d_t S_{\mathbf{n}}(t). \tag{10}$$

It can be shown that the survival probabilities, $S_{\mathbf{n}}$, for all initial states $\mathbf{n} \notin \mathbf{Y}$, satisfy the backward CME,

$$d_t \mathbf{S}(t) = \mathbf{A}_{\mathbf{Y}}^T \mathbf{S}(t), \tag{11}$$

where $\mathbf{S}(t)$ is the vector whose $i^{th}$ component is $S_{\mathbf{i}}(t)$, and $\mathbf{A}_{\mathbf{Y}}$ is the state transition matrix of the CME for the modified system where every state in $\mathbf{Y}$ is absorbing. Taking another time derivative, and using (10), the FPT distributions $\mathbf{f}_{\mathbf{n}}$, for all $\mathbf{n} \notin \mathbf{Y}$, can be seen to also satisfy the backward CME,

$$d_t \mathbf{f}(t) = \mathbf{A}_{\mathbf{Y}}^T \mathbf{f}(t), \tag{12}$$

where $\mathbf{f}(t)$ is the vector whose $i^{th}$ component is $f_{\mathbf{i}}(t)$. The raw moments of the first passage time distribution,

$$\mathbb{E}(\tau_{\mathbf{n}}^k) := \int_0^\infty t^k \mathbf{f}_{\mathbf{n}}(t)\ dt, \tag{13}$$

can be computed by applying the backward master operator $\mathbf{A}_{\mathbf{Y}}^T$ to both sides of (13),

$$\begin{aligned} [(\mathbf{A}_{\mathbf{Y}}^T)^k \mathbb{E}(\boldsymbol{\tau}^k)]_{\mathbf{n}} &= \int_0^\infty t^k [(\mathbf{A}_{\mathbf{Y}}^T)^k \mathbf{f}(t)]_{\mathbf{n}}\ dt \\ &= \int_0^\infty t^k d_t^k f_{\mathbf{n}}(t)\ dt \\ &= k!(-1)^k. \end{aligned} \tag{14}$$

Here we integrated by parts $k$ times in the last step and used the fact that $\int_0^\infty d_t S_{\mathbf{n}}(t)\ dt = -1$. We can write this concisely as,

$$(\mathbf{A}_{\mathbf{Y}}^T)^k \mathbb{E}[\boldsymbol{\tau}^k] = k!(-1)^k \mathbf{1}. \tag{15}$$

#### 3. Analytical solutions of MFPTs for general birth-death process

In this section, we derive analytical solutions of MFPTs for general birth-death processes, where the rates of births and deaths at any given time depends on how many extant molecules there are. This approach uses standard transformation techniques for reducing the order of recurrence relations, and can be found in [4]. We apply this general procedure to derive explicit solutions to three examples of biological relevance: (1) a simple birth-death process, (2) a reduced model of a genetic feedback loop, and (3) a birth-death process with Michaelis-Menten (MM) degradation. Note that examples (1) and (2) appear in the main text, while (3) is used below in Section 4.

*In the following subsections, we will use the simplified notation $\tau_n^N$ to denote the expected waiting time to reach copy number $N$ starting from $n \leq N$, and we will use $\tau_N^n$ to denote the expected waiting time to reach copy number $N$ starting from $n \geq N$. This notation should not be confused with the $N^{th}$ moment of the waiting time distribution.*

##### *3.1. The general procedure*

Consider a general birth-death process with reaction propensities for birth and death given by $a^+(n)$ and $a^-(n)$, respectively. Here $n$ represents the number of molecules in the system. It follows from (15) (or Eq. (7) of the main text) that the expected waiting time to reach $N$ starting from an initial number of molecules $n \leq N$, satisfies the following recurrence relation,

$$(a^+(n) + a^-(n))\tau_n^N - a^-(n)\tau_{n-1}^N - a^+(n)\tau_{n+1}^N = 1, \tag{16}$$

with $\tau_N^N = 0$. There is no need to write special boundary conditions for these equations provided we impose $\tau_n = 0$ for $n < 0$. We now observe that the recurrence equation (16) can be re-expressed in terms of the forward discrete derivative of the sequence $(\tau_n^N)_n$ as follows:

$$a^-(n)(\Delta\tau_{n-1}^N) - a^+(n)(\Delta\tau_n^N) = 1, \tag{17}$$

where $\Delta\tau_i^N$ denotes $\tau_{i+1}^N - \tau_i^N$. As a first order equation, it is convenient to rearrange this as

$$\Delta\tau_n^N = -a^+(n)^{-1}\left(1 - a^-(n)(\Delta\tau_{n-1}^N)\right), \tag{18}$$

where $a^+(n) \neq 0$ for $0 \leq n \leq N$, and we impose $\Delta\tau_n = 0$ for $n < 0$. This is a first-order system and a general formula can be easily extrapolated. Indeed, we can verify by induction that this recursion is solved by,

$$\Delta\tau_n^N = -\sum_{i=0}^{n} \frac{[a^-(n)]^{\underline{i}}}{[a^+(n)]^{\underline{i+1}}}, \tag{19}$$

where for a one-variable function $f(x)$, we let $[f(x)]^{\underline{i}}$ denote $a(x)a(x-1)a(x-2)\cdots a(x-i+1)$. Now using the Fundamental Theorem of (discrete) Calculus, we have that,

$$\sum_{i=n}^{N-1} \Delta\tau_i^N = -\left((\tau_N^N - \tau_{N-1}^N) + (\tau_{N-1}^N - \tau_{N-2}^N) + \cdots + (\tau_{n+1}^N - \tau_n^N)\right) = \tau_N^N - \tau_n^N. \tag{20}$$

Observe that while $N$ appears throughout, we have not yet used $N$ at any point; however as $\tau_N^N = 0$ it follows that

$$\tau_n^N = -\sum_{i=n}^{N-1} \Delta\tau_i^N, \tag{21}$$

and then that

$$\tau_n^N = \tau_0^N - \tau_0^n. \tag{22}$$

Note that (22) should be already expected for a memoryless system, as the time from 0 to $N$ on average should be the time from 0 to $n$ plus the time from $n$ to $N$. Using (21) and the formula in (19) we have that,

$$\tau_n^N = \sum_{i=0}^{N-1}\sum_{j=0}^{i} \frac{[a^-(i)]^{\underline{j}}}{[a^+(i)]^{\underline{j+1}}} - \sum_{i=0}^{n-1}\sum_{j=0}^{i} \frac{[a^-(i)]^{\underline{j}}}{[a^+(i)]^{\underline{j+1}}}. \tag{23}$$

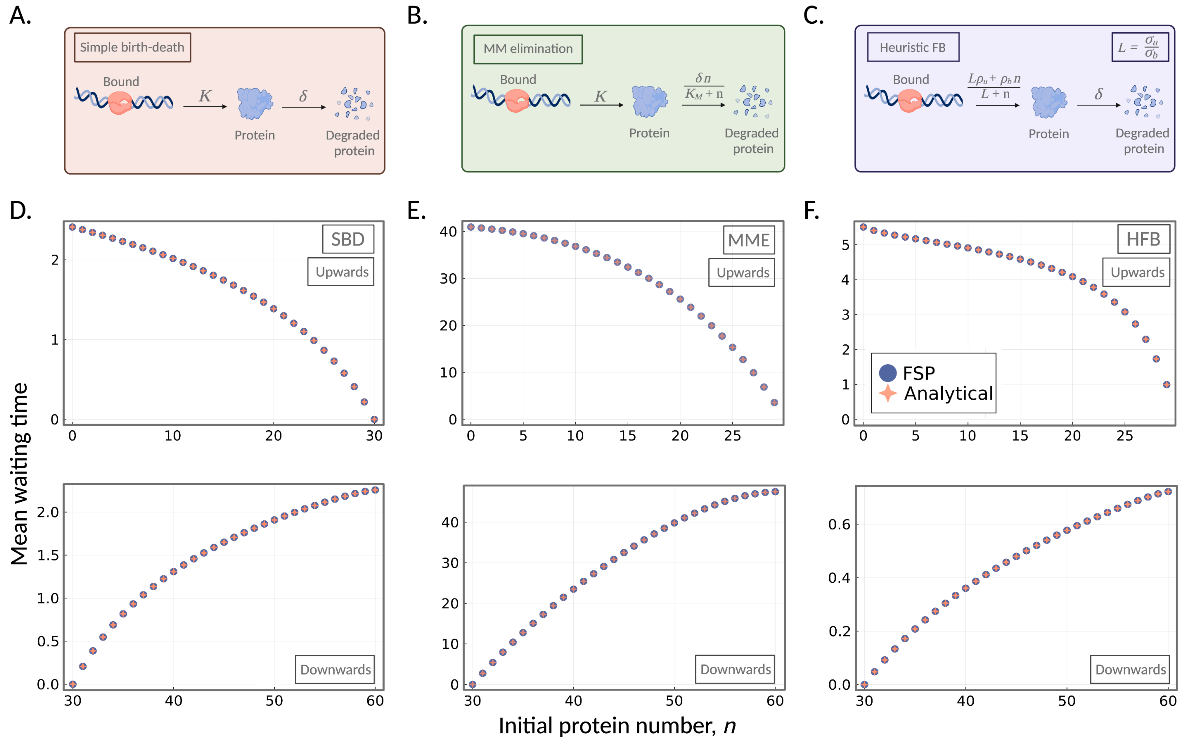


Figure 1: A comparison of derived closed-form solutions for $\tau_n^N$ and $\tau_N^n$ of three birth-death models and the corresponding waiting times obtained from our FSP approach. (A) An illustration of the simple birth-death (SBD) process given in reaction scheme (5). (B) An illustration of the the birth-death process with Michaelis-Menten elimination (MME) given in reaction scheme (32). (C) An illustration of the heuristic feedback model defined in (35). In (D)-(F) (top), we display the mean waiting time of the process to reach a target of $N = 30$ as a function of the initial protein number $n$ ($0 \le n \le N$); this is the upwards case. The analytical solutions (orange) are computed according to (29), (34), and (37), respectively. The mean waiting times computed according to our FSP approach are shown in purple. In (D)-(F) (bottom), we consider the dual case, that is, the mean waiting time of the process to reach a target of $N = 30$ as a function of $n \ge N$; this is the downwards case. The parameters used in (D) for the SBD process are $K = 30$ and $\delta = 1$. For (E) the parameters used for the MME model are $K = 5$, $\delta = 6$ and $K_M = 5$. The parameters of the HFB model in (F) are $\sigma_u = 5$, $\sigma_b$=1, $L = \sigma_u/\sigma_b = 5$, $\rho_u = 10$, $\rho_b = 50$, and $\delta = 2$.

##### *3.2. The dual case*

Assume now that we are interested in the expected waiting time to reach a target $N$ starting from copy number $n \ge N$, which we denote by $\tau_N^n$. As the state-space is now unbounded (or one-sided), we may introduce an absorbing state $J \ge N$ such that no transitions out of $J$ are allowed. This enables us to obtain an approximation to $\tau_N^n$. The recurrence relation for $\tau_N^n$ continues to satisfy (16). Finding a solution for $\tau_N^n$ is essentially the dual problem to solving $\tau_n^N$, and so we can use the general procedure presented above to find a solution for $\tau_N^n$, as we now explain. If we consider $J$ to be the left-hand boundary and $N$ to be the right-hand boundary (as opposed to 0 and $N$ in the case for $\tau_n^N$), we can make the following reinterpretation of the birth and death propensities. Defining $a^+(n) := a^-(J-n)$ and $a^-(n) := a^+(J-n)$, the general solution is now,

$$\tau_N^n = \sum_{i=0}^{J-N-1} \sum_{j=0}^{i} \frac{[a^-(i)]^{\underline{j}}}{[a^+(i)]^{\underline{j+1}}} - \sum_{i=0}^{J-n-1} \sum_{j=0}^{i} \frac{[a^-(i)]^{\underline{j}}}{[a^+(i)]^{\underline{j+1}}}. \tag{24}$$

*Example 1: the simple birth-death process*

Consider again the simple birth-death process introduced above in reaction scheme (5); an illustration of the model can be found in Figure 1(A). Here we provide details of the analytical solution (Eq. (9) of the main text) used in the analysis given in the main text. From (15) (alternatively Eq. (7) of the main text), the expected waiting time to reach $N$ proteins given the system is started from $n \leq N$, can be described by the following recurrence relation,

$$(\delta n + K)\tau_n^N - \delta n \tau_{n-1}^N - K\tau_{n+1}^N = 1, \tag{25}$$

with $\tau_N^N = 0$ and base case at $n = 0$ of $K\tau_0^N - K\tau_1^N = 1$. From (23) it follows that,

$$\tau_n^N = \frac{1}{K}\left(\sum_{i=0}^{N-1}\sum_{j=0}^{i} i^{\underline{j}}\left(\frac{\delta}{K}\right)^j - \sum_{i=0}^{n-1}\sum_{j=0}^{i} i^{\underline{j}}\left(\frac{\delta}{K}\right)^j\right). \tag{26}$$

We may remove the double sum by regrouping as a polynomial in terms of powers of $\frac{\delta}{K}$, which is equivalent to reversing the order of the sums. The coefficient of $\left(\frac{\delta}{K}\right)^m$ in (26) can be seen to be $\sum_{j=0}^{N-1} j^{\underline{m}}$. This is usually written in terms of an equivalent expression involving the sum of rising factorials: $\sum_{j=1}^{N-m} j^{\overline{m}}$, which can be written as,

$$\sum_{j=1}^{N-m} j^{\overline{m}} = \frac{N^{\underline{m+1}}}{m+1}. \tag{27}$$

Thus we have for example,

$$\tau_0^N = \frac{1}{K}\sum_{i=0}^{N-1}\frac{N^{\underline{i+1}}}{i+1}\left(\frac{\delta}{K}\right)^i. \tag{28}$$

The solution for the mean first-passage time of the stochastic system is then given by,

$$\tau_n^N = \frac{1}{K}\left(\sum_{i=0}^{N-1}\frac{N^{\underline{i+1}}}{i+1}\left(\frac{\delta}{K}\right)^i - \sum_{i=0}^{n-1}\frac{n^{\underline{i+1}}}{i+1}\left(\frac{\delta}{K}\right)^i\right). \tag{29}$$

In the main text, we showed how the deterministic waiting time (Eq. (11) of the main text) is a strict upper bound for (29) when $N$ is equal to $\rho K/\delta$ for $0 < \rho < 1$. On the other hand, we can see the deterministic waiting time as the limit, by finding, for any (small) $\varepsilon > 0$ a similar lower bound that converges to,

$$\frac{-\ln(1-(1-\varepsilon)\rho)}{\delta}. \tag{30}$$

To see this, take any positive $0 < \varepsilon < 1$ and consider the truncation of $\mathbb{E}(\tau_0)$ ((29) with $n = 0$) by $i \leq \varepsilon N + 1$. As a truncated sum, this is a strict lower bound for the expected waiting time to $N = \rho K/\delta$ in the stochastic case. Observe that $N^{\underline{i+1}} \geq (N(1-\varepsilon))^{i+1}$, and replacing $N^{\underline{i+1}}$ in the truncated sum by $(N(1-\varepsilon))^{i+1}$ yields the lower bound,

$$\frac{1}{\delta}\sum_{i=0}^{\varepsilon N-1}\frac{((1-\varepsilon)\rho)^{i+1}}{i+1}, \tag{31}$$

which for any fixed $\varepsilon$ (but in the limit of $K/\delta \to \infty$) yields the Taylor series for (30) as claimed.

*Example 2: Michaelis–Menten elimination*

We now consider a birth-death process with Michaelis–Menten elimination. This prototypical reaction network has reactions,

$$\varnothing \underset{\sigma(n)}{\overset{K}{\rightleftharpoons}} P, \tag{32}$$

where $K$ is a constant birth rate and the degradation $\sigma(n)$ is assumed to follow Michaelis-Menten kinetics $\frac{\delta n}{K_M+n}$; here $n$ is the number of protein molecules in the system, $\delta$ is the degradation constant and $K_M$ is the Michaelis-Menten constant. This model has been employed to capture so-called *active protein degradation* where proteins degrade according to some enzymatic reaction, as illustrated in Figure 1(B). Here the birth and death propensities are $a^+(n) = K$ and $a^-(n) = \sigma(n)$, and from (15) (alternatively, Eq. (7) in the main text), the expected waiting time to reach $N$ from $n \leq N$ satisfies the recurrence relation,

$$\left(\frac{\delta n}{(K_M + n)} + K\right)\tau_n^N - \frac{\delta n}{(K_M + n)}\tau_{n-1}^N - K\tau_{n+1}^N = 1, \tag{33}$$

with $\tau_N^N = 0$ and base case at $n = 0$ of $K\tau_0^N - K\tau_1^N = 1$. From (23), we have the following solution for the mean waiting time to reach $N$ from $n \leq N$,

$$\tau_n^N = \frac{1}{K}\left(\sum_{i=0}^{N-1}\sum_{j=0}^{i}\frac{i^{\underline{j}}}{(K_M + i)^{\underline{j}}}\left(\frac{\delta}{K}\right)^j - \sum_{i=0}^{n-1}\sum_{j=0}^{i}\frac{i^{\underline{j}}}{(K_M + i)^{\underline{j}}}\left(\frac{\delta}{K}\right)^j\right). \tag{34}$$

*Example 3: A reduced model of a feedback loop*

Here, we consider one of the most prevalent heuristic stochastic model reduction approaches. As shown in Holehouse and Grima [5], in the limit of fast promotor switching, the stochastic feedback model (reaction scheme given in Eq. (15) of the main text) reduces to just two reactions: an effective zero-order reaction for the production of proteins, and a first-order reaction modelling protein degradation. The effective propensities for the two reactions are defined as follows.

$$a^+(n) := \frac{L\rho_u + \rho_b n}{L + n} \quad \text{and} \quad a^-(n) := \delta n, \tag{35}$$

where $n$ is the number of proteins in the system and $L := \frac{\sigma_u}{\sigma_b}$. Refer to Figure 1(C) for an illustration of the model. From (15) (alternatively Eq. (7) of the main text), we obtain a tridiagonal recurrence relation for the mean waiting time $\tau_n^N$ of the heuristic model,

$$\left(\delta n + \frac{L\rho_u + \rho_b n}{L + n}\right)\tau_n^N - \delta n\tau_{n-1}^N - \left(\frac{L\rho_u + \rho_b n}{L + n}\right)\tau_{n+1}^N = 1, \tag{36}$$

with $\tau_N^N = 0$. From (23), the mean waiting time to reach $N$ starting from $n < N$ is given by,

$$\tau_n^N = \sum_{i=0}^{N-1}\sum_{j=0}^{i}\frac{(L+i)^{\underline{j+1}}}{(L\rho_u + i\rho_b)^{\underline{j+1,\rho_b}}}i^{\underline{j}}\delta^j - \sum_{i=0}^{n-1}\sum_{j=0}^{i}\frac{(L+i)^{\underline{j+1}}}{(L\rho_u + i\rho_b)^{\underline{j+1,\rho_b}}}i^{\underline{j}}\delta^j. \tag{37}$$

Turning to the deterministic regime, the reaction rate equation of the effective model in (35) is given by,

$$\frac{dX}{dt} = \frac{L\rho_u + \rho_b X}{L + X} - \delta X, \tag{38}$$

with $X(0) = 0$. Solving for $t$ gives,

$$t = \frac{1}{\mu^+ - \mu^-}\left[\left(\mu^+ + L\right)\log\left(1 - \frac{X}{\mu^+}\right) - \left(\mu^- + L\right)\log\left(1 - \frac{X}{\mu^-}\right)\right], \tag{39}$$

where $\mu^\pm := \frac{1}{2}\left(\rho_b - L \pm \sqrt{(\rho_b - L)^2 + 4L\rho_u}\right)$. Note that in [5], it has been shown that $\mu^+$ is the steady-state mean of the deterministic system.

In Figure 1 (D)-(F), we verify the accuracy of the solutions given in (29), (34), and (37), and their dual solutions, using comparisons with our FSP approach.

#### 4. Closed-form matrix solutions of FPT problems

In the previous section, we demonstrated how three-term (tridiagonal) recurrence relations arising from a birth-death process can be solved by way of the discrete forward derivative. Here we provide a more general approach that applies to any tridiagonal system of equations. Such systems of equations appear ubiquitously across the sciences, and many approaches have been developed to solve them; see for example [6], and more recently [7]. We illustrate below how our approach can be applied straightforwardly to solve systems with more that one state such as the telegraph model (refer to the reaction scheme given in Eq. (13) of the main text, as well as the surrounding text for details of this model . Consider solving the following general tridiagonal recurrence relation,

$$C_{k-1}x_{k-1} + A_k x_k + B_k x_{k+1} = y_k, \tag{40}$$

for $k \in \{0, 1, 2, \ldots\}$. Rewriting (40) in matrix form, we have the following tridiagonal system of equations,

$$\begin{pmatrix} A_0 & B_0 & & & & & \\ C_0 & A_1 & B_1 & & & & \\ & C_1 & A_2 & B_2 & & & \\ & & C_2 & A_3 & B_3 & & \\ & & & \ddots & & & \\ & & & & & A_{n-1} & B_{n-1} \\ & & & & & C_{n-1} & A_n \end{pmatrix} \begin{pmatrix} x_0 \\ x_1 \\ x_2 \\ x_3 \\ \vdots \\ x_{n-1} \\ x_n \end{pmatrix} = \begin{pmatrix} y_0 \\ y_1 \\ y_2 \\ y_3 \\ \vdots \\ y_{n-1} \\ y_n \end{pmatrix}. \tag{41}$$

Using row operations, we can bring this into upper triangular form,

$$\begin{pmatrix} A'_0 & B_0 & & & & & \\ & A'_1 & B_1 & & & & \\ & & A'_2 & B_2 & & & \\ & & & A'_3 & B_3 & & \\ & & & \ddots & & & \\ & & & & & A'_{n-1} & B_{n-1} \\ & & & & & & A'_n \end{pmatrix} \begin{pmatrix} x_0 \\ x_1 \\ x_2 \\ x_3 \\ \vdots \\ x_{n-1} \\ x_n \end{pmatrix} = \begin{pmatrix} y'_0 \\ y'_1 \\ y'_2 \\ y'_3 \\ \vdots \\ y'_{n-1} \\ y'_n \end{pmatrix}, \tag{42}$$

where the new entries are related to the old ones by the following recursive relations,

$$A'_0 = A_0 \qquad A'_k = A_k - C_{k-1}(A'_{k-1})^{-1}B_k, \qquad (k \geq 1) \tag{43}$$

$$y'_0 = y_0 \qquad y'_k = y_k - C_{k-1}(A'_{k-1})^{-1}y'_k \qquad (k \geq 1). \tag{44}$$

Thus, we have reduced the original second-order system to two simpler recurrence relations of order one. The system can then be solved by back substitution,

$$x_n = (A'_n)^{-1}y_n, \tag{45}$$

$$x_{n-1} = (A'_{n-1})^{-1}(y_{n-1} - B_{n-1}x_n), \tag{46}$$

$$x_{n-2} = (A'_{n-2})^{-1}(y_{n-2} - B_{n-2}x_{n-1}), \tag{47}$$

and so on. As we will see in the following example, there are many cases where the reduced system (45) can be solved straightforwardly by iteration.

##### *4.1. The telegraph model*

The state of the system is given by the number of mRNA and the gene state. We arrange our states as (0, off), (0, on), (1, off), (1, on), (2, off), .... Grouping them in blocks of two we obtain a block tridiagonal system with

$$A_k = \begin{pmatrix} k+\sigma_{\text{on}} & -\sigma_{\text{on}} \\ -\sigma_{\text{off}} & k+\sigma_{\text{off}}+\rho \end{pmatrix}, \tag{48}$$

$$B_k = \begin{pmatrix} & \\ & -\rho \end{pmatrix}, \tag{49}$$

$$C_k = \begin{pmatrix} -k & \\ & -k \end{pmatrix}. \tag{50}$$

$$\tag{51}$$

The entries $A'_k$ satisfy

$$A'_0 = \begin{pmatrix} \sigma_{\text{on}} & -\sigma_{\text{on}} \\ -\sigma_{\text{off}} & \sigma_{\text{off}}+\rho \end{pmatrix}, \tag{52}$$

$$A'_k = \begin{pmatrix} k+\sigma_{\text{on}} & -\sigma_{\text{on}} \\ -\sigma_{\text{off}} & k+\sigma_{\text{off}}+\rho \end{pmatrix} - \rho k (A'_{k-1})^{-1} \begin{pmatrix} & \\ & 1 \end{pmatrix} \qquad (k \geq 1). \tag{53}$$

We can verify by induction that this recursion is solved by

$$A'_k = \begin{pmatrix} k+\sigma_{\text{on}} & -k-\sigma_{\text{on}} \\ -\sigma_{\text{off}} & \sigma_{\text{off}}+\rho \end{pmatrix}. \tag{54}$$

It will be convenient to compute the determinants and inverses of these:

$$d'_k := \det A'_k = \rho(k+\sigma_{\text{on}}), \tag{55}$$

$$(A'_k)^{-1} = \frac{1}{\rho(k+\sigma_{\text{on}})} \begin{pmatrix} \sigma_{\text{off}}+\rho & k+\sigma_{\text{on}} \\ \sigma_{\text{off}} & k+\sigma_{\text{on}} \end{pmatrix}. \tag{56}$$

The right-hand side of the MFPT equation is changed to

$$y'_k = \left( \sum_{i=0}^{k} k(k-1)\dots(k-i)(A'_{k-i}A'_{k-i+1}\dots A'_{k-1})^{-1} \right) \begin{pmatrix} 1 \\ 1 \end{pmatrix}. \tag{57}$$

Our system of equations is thus

$$\begin{pmatrix} A'_0 & \begin{smallmatrix} 0 & 0 \\ 0 & -\rho \end{smallmatrix} & & & & & \\ & A'_1 & \begin{smallmatrix} 0 & 0 \\ 0 & -\rho \end{smallmatrix} & & & & \\ & & A'_2 & \begin{smallmatrix} 0 & 0 \\ 0 & -\rho \end{smallmatrix} & & & \\ & & & A'_3 & \begin{smallmatrix} 0 & 0 \\ 0 & -\rho \end{smallmatrix} & & \\ & & & \ddots & & & \\ & & & & & A'_{n-1} & \begin{smallmatrix} 0 & 0 \\ 0 & -\rho \end{smallmatrix} \\ & & & & & & A'_n \end{pmatrix} \begin{pmatrix} x_0 \\ x_1 \\ x_2 \\ x_3 \\ \vdots \\ x_{n-1} \\ x_n \end{pmatrix} = \begin{pmatrix} y'_0 \\ y'_1 \\ y'_2 \\ y'_3 \\ \vdots \\ y'_{n-1} \\ y'_n \end{pmatrix}. \tag{58}$$

Since we have the inverses of the $A'_k$, this can be computed straightforwardly using backsubstitution.

#### 5. Computational efficiency of the modified FSP

In this section, we present three applications of our FSP approach for computing mean first-passage times (MFPTs). We demonstrate the versatility of the approach by applying the method to three exemplary models from the literature: (1) a birth-death process with Michaelis-Menten degradation; (2) a compartmental model of disease spread; and, (3) a Michaelis-Menten reaction scheme. Our approach is shown to be significantly more computationally efficient than traditional Monte-Carlo simulations. All simulations were run on a 2.6 GHz 6-Core Intel Core i7 processor with 32GB RAM.

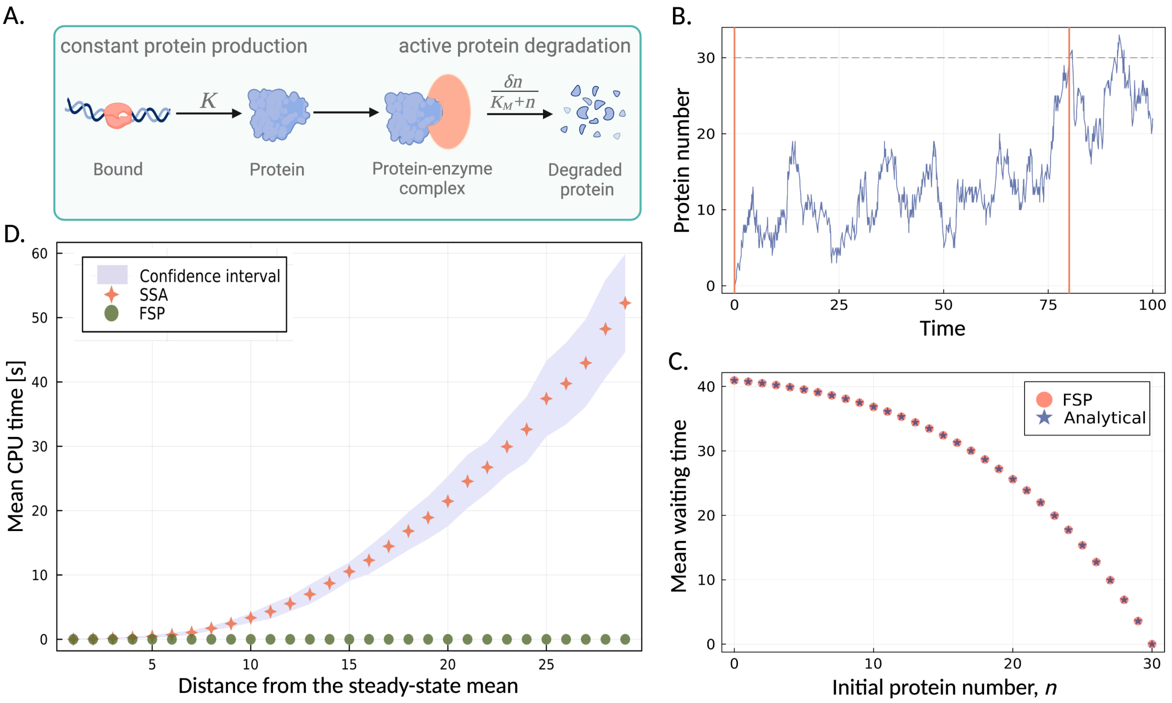


Figure 2: Computational cost of the FSP approach to compute the MFPT distribution using a one dimensional birth-death system with Michaelis–Menten (MM) degradation kinetics. (A) An illustration of a birth-death system with MM degradation kinetics. Protein is produced constantly at rate $K$ and actively degraded by an enzymatic reaction with rate $\delta n/(K_M + n)$. (B) The protein number as a function of time. The system is initialised at $n = 0$ (first orange vertical line) and stopped when $n$ first reaches the threshold value of $N = 30$. The grey dotted line represents the threshold value and the second orange vertical line indicates the first time the system reaches the threshold value. (C). The mean waiting time as a function of the initial protein number. (D) Comparison of the CPU time taken to calculate the MFPT distribution using FSP (orange) and the SSA (green) as a function of the initial protein number. The purple shading indicates the 95% confidence interval. The model parameters used are: $K_b = 5$; $K_d = 6$; $K_M = 5$. The SSA is run for the number of trajectories it takes for the MFPT to agree with the MFPT given by the FSP, with an error tolerance of 10%.

*Example 1: A birth-death model with Michaelis–Menten degradation*

Consider again the simple birth-death system with Michaelis–Menten degradation given in (32) above (see Figure 2(A)). We are interested in the time that it takes for the protein number $n$ to reach a certain threshold value $N$. Here, we set $N$ to be the steady-state mean of the system, and consider the mean waiting time of the system to reach this threshold, conditional on the system starting from some initial state $\mathbf{n}_0$. This information provides insights into the underlying biochemical processes and the timescales associated with protein-enzyme interactions.

In Figure 2(B), we display a representative time series of the protein number $n$. The vertical orange lines indicate the times at which the protein count is $n = 0$ and when it has reached the steady-state mean ($N = 30$). From the single trajectory presented, we observe that it takes approximately 78 time units for the protein count to first reach the threshold $N = 30$, given the number of proteins in system is initially $n = 0$. As this is a stochastic process, the mean of the first passage time is a more representative measure of the process dynamics. One advantage of our approach for computing mean first passage times is that it allows us to simultaneously determine the MFPT for all initial protein counts that are less than the threshold value. In Figure 2(C), we plot the mean waiting time to reach the steady-state value ($N = 30$) for different initial protein numbers. Here we let the initial protein $\mathbf{n}$ vary between 0 and 30. As expected, the lower the initial protein count, the longer the system takes to reach the threshold value.

We next compare the computational time of computing MFPTs using our FSP approach with that of the stochastic simulation algorithm (SSA). Figure 2(D) shows the mean processing time required to obtain the

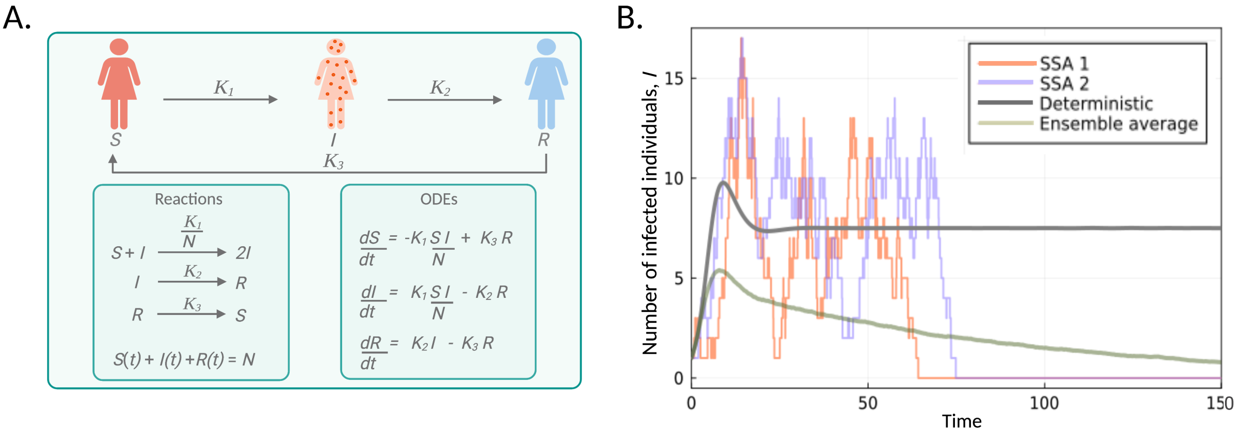


Figure 3: The deterministic and stochastic dynamics of an epidemic SIRS model. (A) An illustration of the SIRS model where the population is divided into susceptible (S), infected (I) and recovered (R) individuals. The typical trajectory is for an individual to move from S to I to R, where individuals in state R can again move to state S due to a waning immunity. The chemical reactions are given in the left-hand side box and the associated reaction rate equations are shown in the right-hand side box. (B) The deterministic model (grey curve) predicts that the number of infected individuals in a population reaches a non-zero steady-state. Stochastic trajectories obtained using the SSA (orange and purple curves), however, are driven to the extinction state by random fluctuations. The ensemble average of 5 x $10^3$ stochastic trajectories is shown in the green curve and differs distinctly from the deterministic prediction.

MFPT conditional on the initial protein number, using the stochastic simulation algorithm (shown in orange) and our FSP approach (shown in green). The computation time for the MFPTs shown in Figure 2(C) is on average 0.26 milliseconds when using the FSP approach, and is approximately 53 seconds when using the SSA. This difference in computation time is due to the fact that the FSP approach allows for the calculation of the MFPT for *all* initial states $\mathbf{n}$ in a single computation, whereas the SSA requires the computation of multiple trajectories for each initial condition $\mathbf{n}$ in order to construct the MFPT curve given in Figure 2(C).

*Example 2: An epidemic SIRS model*

We here consider one of the simplest compartmental models in epidemiology: the SIRS model [8, 9]. Here the population is partitioned into three compartments: Susceptible ($S$), Infected ($I$), and Recovered ($R$) individuals. The total population size, $N$, is fixed such that $S + I + R = N$. Individuals in state $S$ are considered to be at risk of contracting the disease, but are currently uninfected. Individuals in state $I$ have been infected with the disease and are capable of transmitting it to others. Individuals in state $R$ are recovered, but are able to become susceptible again – modelling a waning immune response.

The SIRS model is defined by the following reaction equations,

$$S + I \xrightarrow{K_1/N} 2I, I \xrightarrow{K_2} R, R \xrightarrow{K_3} I, \tag{59}$$

where $K_1/N$ is the transmission rate, $K_2$ is the rate of recovery and $K_3$ is the reinfection rate. An illustration of the SIRS model is shown in Figure 3(A). The basic reproduction number, $\mathfrak{R}_0$, is equal to $K_1/K_2$; this ratio is derived as the expected number of new infections from a single infection in a population where all subjects are susceptible. Note that when solving (59), we exploit the conservation law $S = N - R - I$, giving us an effective two-compartmental model in $I$ and $R$. The associated deterministic equations of the two-compartmental model are,

$$\frac{dI}{dt} = \frac{K_1(N - I - R)I}{N} - K_2 I, \tag{60}$$

$$\frac{dR}{dt} = K_2 I - K_3 R. \tag{61}$$

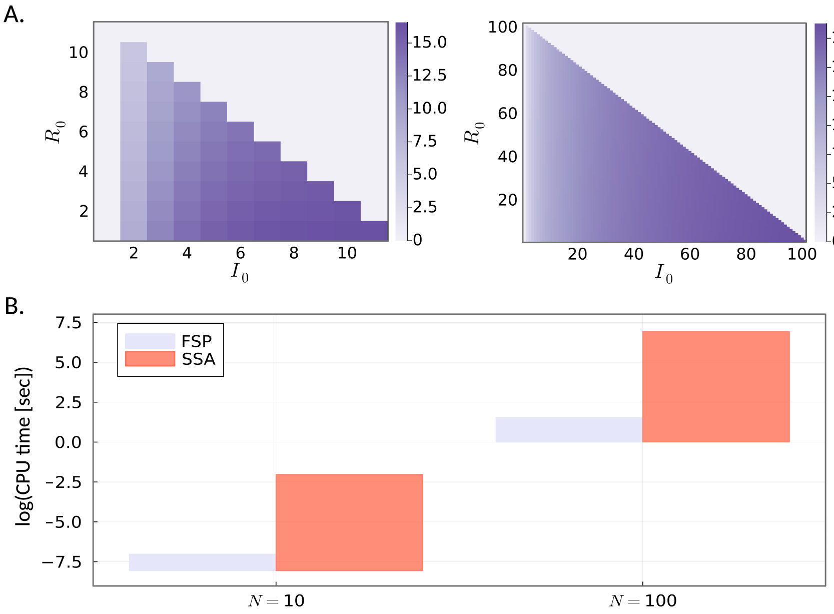


Figure 4: Computational cost of the FSP approach to compute the MFPT distribution using an epidemic SIRS model of infection. (A) Heatmaps showing the mean time to extinction as a function of the parameters $I_0$, the initial number of infected individuals, and $R_0$, the initial number of recovered individuals, for a population size of $N = 10$ (left panel) and $N = 100$ (right panel). (B) Comparison of the median CPU time of the FSP approach (purple) and the SSA (orange) as a function of the population size $N$. Note that for $N = 10$, we re-normalise the baseline CPU time for visualisation purposes. Parameters used: $K_1 = 0.5$, $K_2 = 0.5$, $K_3 = 0.3$, $N = 10$ (left panel). $K_1 = 0.05$, $K_2 = 0.3$, $K_3 = 0.3$, $N = 100$ (right panel).

When the basic reproduction number $\mathcal{R}_0 > 1$, the deterministic regime realises a so-called *endemic equilibrium* (a non-zero steady-state), and, as such, predicts that the disease will never die out. In comparison, the stochastic regime is able to converge to the disease-free state. We illustrate this behaviour in Figure 3(B), where we plot the number of infected individuals over time. Trajectories of the SIRS model simulated using the SSA fluctuate around the deterministic steady-state mean for a period of time, before fluctuations drive the trajectories to zero (orange and blue trajectories). The stochastic ensemble average (green curve) converges to zero and thus disagrees with the deterministic prediction (grey curve). As such, it is necessary to turn to a stochastic framework when modelling the extinction time of an infectious disease. Using a stochastic framework, we can model the duration of the epidemic as a first passage time problem.

We define the extinction time as the expected waiting time of the system in (59) to reach zero infected individuals, given it was initialised with a pre-specified number of infected ($I_0$), susceptible ($S_0 = N - I_0 - R_0$), and recovered ($R_0$) individuals. We are interested in computing the mean time to extinction only in the stochastic regime since the deterministic regime cannot achieve disease extinction. In Figure 4(A) we plot heatmaps of the extinction time, as a function of the initial number of infected $I_0$ and recovered $R_0$ individuals, for population size $N = 10$ and $N = 100$, using our FSP approach. The extinction time is greatly affected by the initial number of infected individuals. The larger the proportion of infected individuals is at the start of the epidemic, the longer it takes the epidemic to die out, and vice-versa. It is well-appreciated that the time to extinction, in SIR(S) models, is profoundly affected by the both the initial number of infected individuals and the size of the population [10].

Next, we are interested in the average CPU time taken to achieve the results in Figure 4(A), using our FSP approach and the SSA. Figure 4(B) shows the median CPU time in seconds for $N = 10$ and $N = 100$ for both the FSP (orange) and SSA (purple). The FSP is considerably more time efficient than the SSA, approximately four orders of magnitude when $N = 10$ and seven orders of magnitude when $N = 100$. Reasoning for the reduction in computational time follows the same logic as the birth-death model with MM degradation.

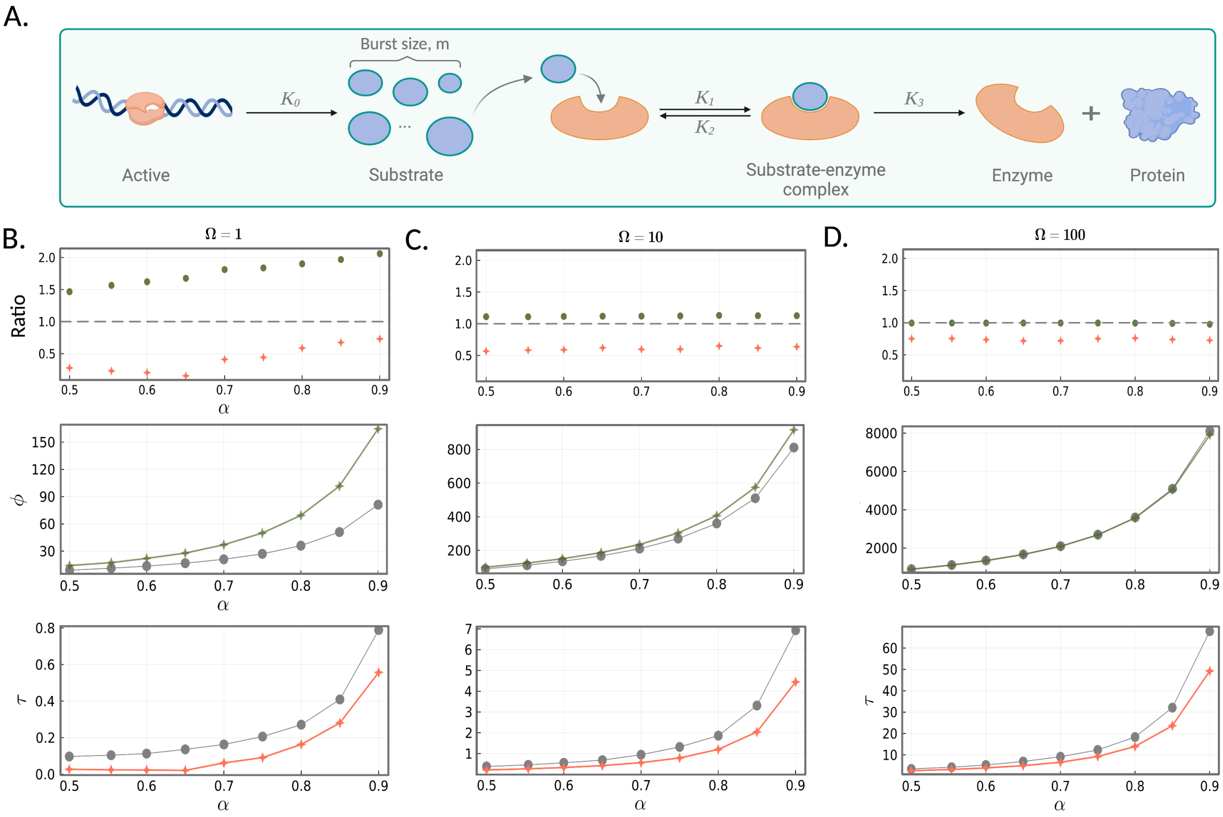


Figure 5: The Michaelis-Menten equation in steady-state conditions. (A) An illustration of the Michaelis-Menten reaction mechanism given by the reaction scheme in (62). In (B)-(D) (top panels) we plot the substrate ratios $\varphi_{\text{sto}}/\varphi_{\text{det}}$ (green circles) and the associated waiting time ratios $\tau_{\text{sto}}/\tau_{\text{det}}$ (orange stars), as a function of the ratio $\alpha$ (a measure of the enzyme saturation). In the middle panels, we compare $\varphi$ under deterministic (grey) and stochastic (green) models. In the bottom panels we compare $\tau$ under deterministic (grey) and stochastic (orange) models, as a function of $\alpha$. (B) When the volume $\Omega$ is 1, the discrepancy between $\varphi$ and $\tau$ is largest, and increases as $\alpha$ approaches 1. As the volume $\Omega$ increases, from (C) 10 to (D) 100, the discrepancy decreases. Model parameters used: $K_1 = 4$, $K_2 = 3$, $K_3 = 37$, $E_T = 60$ and $\Omega \in \{1, 10, 100\}$. $\alpha$ is varied through $K_0$.

*Example 3: Michaelis-Menten kinetics in steady-state conditions*

We consider a generalised stochastic model of the well-studied Michaelis-Menten reaction mechanism [11], where the enzyme kinetics are confined to a sub-cellular compartment of volume $\Omega$. The reactions are given by,

$$\varnothing \xrightarrow{K_0} mS, \quad S + E \underset{K_2}{\overset{K_1}{\rightleftharpoons}} ES \xrightarrow{K_3} E + P. \tag{62}$$

Here, a substrate molecule $S$ is spontaneously created in bursts and continuously supplied to the sub-cellular compartment at rate $K_0$. In the sub-cellular compartment, $S$ reversibly binds to a free enzyme molecule $E$ with rates $K_1$ and $K_2$, respectively, to form the complex $ES$. The complex subsequently yields a product molecule $P$, with rate $K_3$. The parameter $m$ represents the burst size, which for simplicity, we take here to be a constant (see Figure 5(A)). The total number of enzyme molecules $E_T$ is conserved such that $E_T = E + ES$. The system has a steady state in $S$ if $\alpha = \frac{mK_0\Omega}{K_3 E_T} \leq 1$, in other words, if the supply rate of $S$ is less than the maximum production rate of $P$. In this way, the ratio $\alpha$ provides a measure of how saturated the enzyme is with substrate. Note that $\alpha > 1$ implies that the substrate numbers increase unboundedly with time.

We assess the validity of the deterministic description of the enzyme kinetics by comparing to exact stochastic simulations of the MM reaction scheme ((62)). Specifically, we are interested in the steady-state concentration of $S$, and the associated mean waiting time to reach the steady-state, given that the system is started with zero substrate and all enzyme initially in the unbound, free state. Let $\varphi$ be 90% of the steady-state mean substrate concentration and $\tau$ be the expected waiting time of the system to reach $\varphi$. To obtain $\varphi$ in the deterministic regime, we solve the deterministic reaction rate equations at steady-state. The associated expected waiting time $\tau$ is computed by solving the rate equations and recording the first time that $S$ reaches $\varphi$. In the stochastic regime, we obtain $\varphi$ by way of SSA. We take the average over 1000 independent simulations of (62) for a time-span of $10^3$, collecting every 20 time units. We utilise our FSP approach to compute the associated mean waiting time, $\tau$. All simulations are performed in the Julia Programming Language [12].

In Figure 5(B)-(D), we examine the effect of a sub-cellular compartmental volume on both $\varphi$ and the associated mean waiting time $\tau$, between the stochastic and deterministic regimes. The two regimes are compared for increasing volumes, $\Omega \in \{1, 10, 100\}$. In the top panels, we consider the substrate ratio, given as $\varphi_{\text{sto}}/\varphi_{\text{det}}$, and the expected-waiting-time ratio, given as $\tau_{\text{sto}}/\tau_{\text{det}}$, as a function of the saturation constant $\alpha$. Note that when the deterministic and stochastic regimes are in agreement, the value of these ratios is unity. In the low molecular regime (B) & (C), the stochastic simulations consistently yield higher substrate concentrations than those predicted by deterministic rate equations. This discrepancy becomes more significant as $\alpha$ increases, and can also be observed in the remaining plots, which illustrate the dependence of $\varphi$ (middle panel) and $\tau$ (bottom panel) on $\alpha$. As $\alpha$ approaches unity, the system becomes unstable, and the fluctuations around the mean substrate concentration become increasingly large, causing significantly higher $\varphi$ values in the stochastic regime. This results in an infinite ratio between the two regimes in the limit. Conversely, the waiting time for the system to reach $\varphi$ is consistently lower in the stochastic regime, despite higher substrate concentrations. This can be seen in the ratio plots (top panel), where the mean waiting time ratio is always below one, despite the substrate ratio consistently being greater than or equal to one. The bottom panel plots clearly show that the waiting time $\tau$ in the stochastic regime (orange stars) is always less than the deterministic regime (grey dots). Thus, increasing molecular noise in the system (by decreasing the system size $\Omega$) decreases the waiting time to reach $\varphi$, implying that intrinsic noise profoundly affects the mean first-passage time. For larger systems (Figure 5(D)), we observe convergence between the stochastic and deterministic substrate concentrations. However, we can see that the waiting times converge significantly more slowly, requiring a much larger system size for convergence.

#### 6. Parameters for Figure 4 of main text

Table 1: Model parameters used in Figure 4 of the main text

| **Fig. 4.** | **Panel** | **Parameters** | | | | **Mean** | |
|---|---|---|---|---|---|---|---|
| | | $\sigma_b$ | $\sigma_u$ | $\rho_b$ | $\rho_u$ | Deterministic | Stochastic |
| (B) | Top and middle | 0.03 | 60 | 300–6500 | 10 | 11.68–4504.44 | 11.68–4487.80 |
| | Bottom | 0.03 | 0.1–60 | 500–6500 | 10 | | |
| (C) | Top and middle | 10 | 10 | 300–6500 | 10 | 300–6499.01 | 299.07–6498.60 |
| | Bottom | 10 | 0.1–6 | 300-6500 | 10 | | |
| (D) | Top and middle | 0.04 | 0.1 | 10–3500 | 3500 | 97.36–3500 | 44.36–3499.90 |
| | Bottom | 0.04 | 0.1–10 | 10–1000 | 1000 | | |
| (E) | Top and middle | 10 | 10 | 100–3500 | 3500 | 126.64–3500.64 | 126.24–3500 |
| | Bottom | 10 | 0.1–10 | 100–1000 | 1000 | | |
